# New design strategies for ultra-specific CRISPR-Cas13a-based RNA-diagnostic tools with single-nucleotide mismatch sensitivity

**DOI:** 10.1101/2023.07.26.550755

**Authors:** Adrian M. Molina Vargas, Raven Osborn, Souvik Sinha, Pablo R. Arantes, Amun Patel, Stephen Dewhurst, Giulia Palermo, Mitchell R. O’Connell

## Abstract

The pressing need for clinical diagnostics has required the development of novel nucleic acid-based detection technologies that are sensitive, fast, and inexpensive, and that can be deployed at point-of-care. Recently, the RNA-guided ribonuclease CRISPR-Cas13 has been successfully harnessed for such purposes. However, developing assays for detection of genetic variability, for example single-nucleotide polymorphisms, is still challenging and previously described design strategies are not always generalizable. Here, we expanded our characterization of LbuCas13a RNA-detection specificity by performing a combination of experimental RNA mismatch tolerance profiling, molecular dynamics simulations, protein, and crRNA engineering. We found certain positions in the crRNA-target-RNA duplex that are particularly sensitive to mismatches and establish the effect of RNA concentration in mismatch tolerance. Additionally, we determined that shortening the crRNA spacer or modifying the direct repeat of the crRNA leads to stricter specificities. Furthermore, we harnessed our understanding of LbuCas13a allosteric activation pathways through molecular dynamics and structure-guided engineering to develop novel Cas13a variants that display increased sensitivities to single-nucleotide mismatches. We deployed these Cas13a variants and crRNA design strategies to achieve superior discrimination of SARS-CoV-2 strains compared to wild-type LbuCas13a. Together, our work provides new design criteria and new Cas13a variants for easier-to-implement Cas13-based diagnostics.

**KEY POINTS:**

- Certain positions in the Cas13a crRNA-target-RNA duplex are particularly sensitive to mismatches.
- Understanding Cas13a’s allosteric activation pathway allowed us to develop novel high-fidelity Cas13a variants.
- These Cas13a variants and crRNA design strategies achieve superior discrimination of SARS-CoV-2 strains.

**GRAPHICAL ABSTRACT:** New strategies to improve Cas13a RNA-detection specificity developed via mismatch tolerance profiling, uncovering features that modulate specificity, and structure-guided engineering of LbuCas13a.

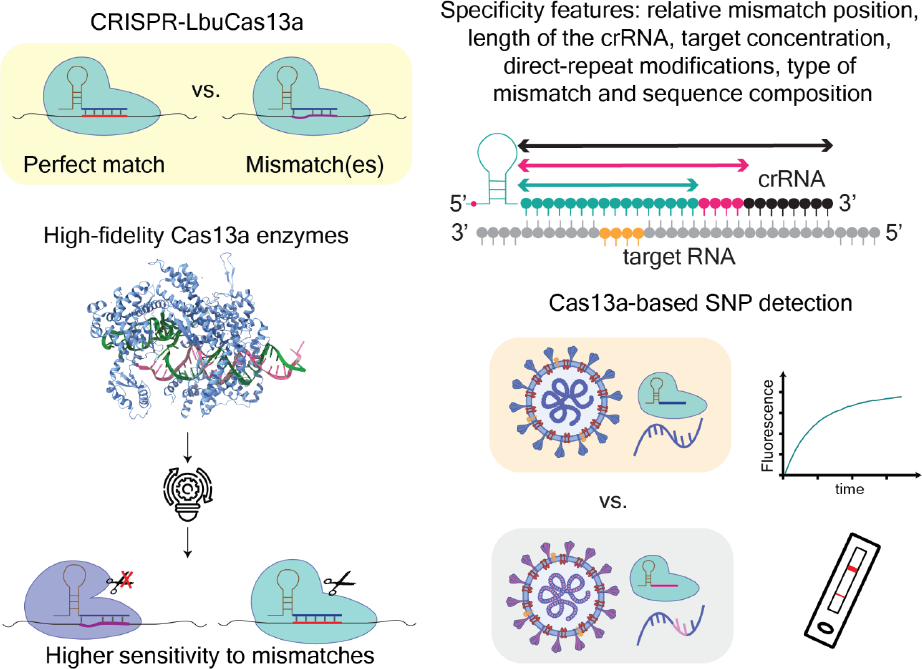

## INTRODUCTION

RNA-guided nucleases use base-pair complementarity between small guide RNAs and target DNA or RNA for specific gene regulation and/or defense against foreign nucleic acids across different organisms. In particular, CRISPR (Clustered Regularly Interspaced Short Palindromic Repeats) and their associated (Cas) protein effectors have evolved as powerful immune systems that protect prokaryotes against mobile genetic elements (1). CRISPR-Cas systems are incredibly diverse and are classified in various types (I-VI) (1–4) and a select group of them can target and degrade RNA (2). Of interest here, Type VI CRISPR-Cas systems contain a single effector ribonuclease, Cas13, that binds and processes a CRISPR-RNA (crRNA), forming an RNA-guided RNA-targeting complex. This effector complex, upon finding sufficient base-pair complementarity with a target RNA, is able to engage in single stranded RNA (ssRNA) cleavage activity mediated by its two Higher Eukaryotes and Prokaryotes Nucleotide (HEPN) domains (reviewed in (5)). Because of its programmable and versatile nature, Cas13 holds promise as a powerful addition to the RNA toolkit. Consequently, Cas13 has been successfully engineered for potent RNA-knockdown in eukaryotic cells with minimal off-target effects, and other exciting applications described to date include utilizing a nuclease-dead Cas13 (dCas13) as a programmable RNA-binding protein for RNA imaging, RNA-splicing, RNA-detection or RNA-editing applications, among others (6–13).

Once active, Cas13 degrades not only its bound target RNA (*cis*-cleavage) but also other ssRNAs present, in a non-specific manner. This *trans*- or collateral-cleavage activity has been harnessed for the development of ultrasensitive RNA detection tools. Detection occurs when Cas13 and a programmed crRNA recognizes the RNA of interest and cleaves an RNA reporter which can be measured by fluorescence (14,15), lateral flow detection or other modalities such electrochemical biosensors, depending on the design of the reporter (16–20). The same principle of detecting cleaved reporters has been harnessed for other CRISPR and/or RNA-guided nucleases such as CRISPR-Cas12 or prokaryotic Argonaute proteins (16,21–30).

For Cas13, multiple efforts in recent years have focused on establishing powerful platforms that couple nucleic acid amplification to Cas13-detection for higher sensitivity (SHERLOCK, CREST), streamline single-step protocols (SHINE), perform multiplex testing (CARMEN) or use different Cas13 orthologs (SENSR) (16,17,31–35). These robust platforms are promising to be deployed at point-of-care locations making testing accessible, scalable, faster, low-cost, and flexible, which is particularly relevant in the context of fast-developing outbreaks. As a result, Cas13 has been shown to be a robust option to detect SARS-CoV-2 in patient samples and currently some Cas13-based assays have received FDA Emergency Use Authorization (36,37).

However, exploration of the principles for crRNA design in the context of Cas13-based diagnostics has been limited, requiring a case-by-case design process to identify crRNAs that provide high activity with low off-target activation of Cas13. This is particularly relevant when trying to distinguish genetic variation within a sample containing potentially containing a pathogen or disease variant of interest. For example, previous studies with SHERLOCK and SATORI (CRISPR-based amplification-free digital RNA detection) (16,17,38) showed that in most cases LwaCas13a is resistant to single nucleotide mismatches, which could be advantageous to detect rapidly changing pathogens but also poses a challenge when discrimination of single-nucleotide polymorphisms (SNP) is desirable. One successful strategy has been to introduce one or more additional ‘synthetic mismatches’ to achieve discrimination at a critical position (16,17,28,39). However, where to place the discriminatory and/or synthetic mismatches is not intuitive, needs to be determined empirically for each crRNA-target set and does not always yield robust discrimination (29,32,40). Recent work has exploited machine learning principles to discern nucleotide-preferences, mismatch tolerance and design crRNAs for viral detection that are efficient when testing or discriminating genetic variation between samples (ADAPT) (41), but still require thorough validation. Therefore, thus more work in generating prediction tools that can design crRNAs with maximal single nucleotide polymorphism (SNP) discrimination, and/or new crRNA design strategies or highly specific CRISPR enzymes that can differentiate closely related sequences is required to overcome this challenge. Further progress in the diagnostic field will require a deeper understanding of the biophysical parameters that underlie Cas13 RNA-recognition and activation, which in turn will guide the rational design of more specific Cas13 RNA-diagnostics.

Previous work suggests that Cas13, like other RNA-guided enzymes and CRISPR-Cas effectors, has a differential sensitivity to mismatches depending where the mismatch occurs in the crRNA-spacer: target region (8–10,42,43). Additionally, there is precedent that the length of the spacer plays a role in activity and specificity of Cas13 (8,10,43,44). To further ascertain the contributions of each of these features, we selected one Cas13 ortholog that has been extensively used for a number of applications, LbuCas13a. Tambe *et al.* had previously established fundamental regions in the crRNA-target duplex that gate binding and nuclease activation in LbuCas13a (45). Here we expand upon these foundational studies by further exploring how new target sequence contexts (e.g., GC content), the contributions of crRNA length and changes in the crRNA architecture affect LbuCas13’s sequence specificity, and specifically tolerance to mismatches. We find that much like *Tambe et al.,* LbuCas13a’s sensitivity to mismatches is crRNA length and position-dependent and identified sensitive positions that prevent LbuCas13a activation if a mismatch occurs. Additionally, we recently we used molecular dynamic simulations to understand allostery and as a result identified several mutations that alter allosteric communication pathways in LbuCas13a (*Sinha et al.*, manuscript under review). Here we further explored these variants and observed that they are also more sensitive to mismatches experimentally, as well as exhibit altered allosteric communication pathways relative to WT LbuCas13a. We harnessed these sensitivities in combination to our newly described LbuCas13a variants, yielding highly specific discrimination of RNA-targets down to single-nucleotide polymorphisms (SNP). We deployed this novel platform for the detection of single-nucleotide polymorphisms in SARS-CoV-2 variants of concern and showed their potential for disease diagnostics. Finally, in the context of a Cas13a targeting reaction, we show that the orientation in the base pairs participating in a crRNA:target-RNA mismatch, and the sequence context around this mismatch, are features that modulate the tolerance for this mismatch, and this additional design complexity needs to be considered when designing crRNAs for Cas13-based diagnostics.

## MATERIAL AND METHODS

### Cas13a protein expression and purification

For expression of wild-type LbuCas13a we used a plasmid that contains a codon-optimized Cas13a sequence which is N-terminally tagged with a His6-MBP-TEV-protease cleavage site sequence. (Addgene Plasmid #83482, East-Seletsky et al. (15)). LbuCas13a variants were generated from the wild-type vector via site-directed mutagenesis using the primers indicated in **Table S1**.

Purification of all constructs was carried out as previously described, with some modifications (14,15). Briefly, expression vectors were transformed into Rosetta2 DE3 grown in LB media supplemented with 0.5% w/v glucose at 37 °C. Protein expression was induced at mid-log phase (OD_600_ ∼0.6) with 0.5 mM IPTG, followed by incubation at 16 °C overnight. Cell pellets were resuspended in lysis buffer (50 mM HEPES [pH 7.0], 1 M NaCl, 5 mM imidazole, 5% (v/v) glycerol, 1 mM DTT, 0.5 mM PMSF, EDTA-free protease inhibitor [Roche]), lysed by sonication, and clarified by centrifugation at 15,000g. Soluble His_6_-MBP-TEV-Cas13a was isolated over metal ion affinity chromatography, and in order to cleave off the His_6_-MBP tag, the protein-containing eluate was incubated with TEV protease at 4 °C overnight while dialyzing into ion exchange buffer (50 mM HEPES [pH 7.0], 250 mM NaCl, 5% (v/v) glycerol, 1 mM DTT). Cleaved protein was loaded onto a HiTrap SP column (GE Healthcare) and eluted over a linear KCl (0.25–1 M) gradient. LbuCas13a containing fractions were pooled, concentrated, and further purified via size-exclusion chromatography on a S200 column (GE Healthcare) in gel filtration buffer (20 mM HEPES [pH 7.0], 200 mM KCl, 5% glycerol (v/v), 1 mM DTT), snap-frozen in liquid N_2_ and were subsequently stored at −80°C.

### *In-vitro* RNA transcription

Mature crRNAs were synthetically made by IDT. All RNA targets were transcribed *in vitro* using previously described methods (15,46). Briefly, all targets were transcribed off a single-stranded DNA oligonucleotide template (IDT) using T7 polymerase. were annealed to a 1.5-fold molar excess of an oligonucleotide corresponding to the T7 promoter sequence (5′-GGCGTAATACGACTCACTATAGG-3′). Transcription reactions were incubated at 37°C for 3 h and contained 1 μM template DNA, 100 μg/mL T7 polymerase, 1 μg/mL pyrophosphatase (Roche), 5 mM NTPs, 30 mM Tris-Cl (pH_RT_ 8.1), 25 mM MgCl_2_, 10 mM dithiothreitol (DTT), 2 mM spermidine, and 0.01% Triton X-100. Reactions were then treated with 5 units of DNase (Promega) and incubated for an additional 30 min at 37°C before being loaded on a 15% urea-polyacrylamide gel. Transcribed RNAs were purified using 15% Urea-PAGE. RNAs were excised from the gel and eluted into DEPC water overnight at 4°C followed by ethanol precipitation. RNAs were resuspended in DEPC water and stored at −80°C. All sequences can be found in **Table S2**.

### Fluorescent ssRNA nuclease assays

Cas13 trans-cleavage nuclease activity assays were performed as previously described with some modifications (East-Seletsky et al., 2017). Briefly, 100 nM LbuCas13a:crRNA complexes were assembled in cleavage buffer (20 mM HEPES-Na pH 6.8, 50 mM KCl, 5 mM MgCl_2_, 10 μg/mL BSA, 100 μg/mL tRNA, 0.01% Igepal CA-630 and 5% glycerol), for 30 min 37°C. 100 nM of RNase Alert reporter (IDT) and various final concentrations of ssRNA-target were added to initiate the reaction. These reactions were incubated in a fluorescence plate reader (Tecan Spark) for up to 120 min at 37°C with fluorescence measurements taken every 5 min (λ_ex_: 485 nm; λ_em_: 535 nm). Time-course and end-point values at 1 hour were background-subtracted, normalized, and analyzed with their associated standard errors using Prism9 (GraphPad) and R version 4.1.1. Target RNAs and crRNAs used in the study can be found in **Table S3**.

For lateral flow based detection, we generated the reaction mix as described above, except we used a biotinylated FAM reporter at a final concentration of 1 µM rather than the RNAse Alert substrate. After 30 minutes of incubation at 37 °C, the detection reaction was diluted 1:4 in Milenia HybriDetect Assay Buffer, and the Milenia HybriDetect 1 (TwistDx) lateral flow strip was added. Sample images were collected 5 min following incubation of the strip.

### Molecular Dynamics simulations

Molecular dynamics (MD) simulations were based on the structure of the *Leptotrichia buccalis* (Lbu) Cas13a bound to a crRNA and a tgRNA (PDB: 5XWP), obtained by single-wavelength anomalous diffraction at 3.08 Å resolution (47). Four Cas13a: crRNA: target-RNA complexes were considered, including either a perfectly matched crRNA: target-RNA duplex, or crRNA: target-RNA duplexes that contain a single mismatch at either spacer nucleotide position 4, 7, or 11. The sequences of spacer crRNA and target RNAs (tgRNA) that are used for the complexes are the same as used for their corresponding cleavage assays. In all systems, we reinstated the catalytic H1053 and R1048 in the HEPN domains, which were mutated in alanine in the experimental structures (same reference). The systems were solvated, leading to simulation cells of ∼ 146 * 95 * 142 Å^3^, and neutralized by the addition of an adequate number of Na^+^ ions.

MD simulations were performed by employing a simulation protocol tailored for protein/nucleic acid complexes, previously employed for CRISPR-Cas systems (48–52), and also used in our companion paper. We employed the Amber ff19SB force field (53) with the *χOL3* corrections for RNA (54,55). The TIP3P model was used for explicit water molecules (56). As a first step, all systems were subjected to energy minimization to overcome the potential inter-and intra-molecular steric clashes. Then, the systems were heated from 0 to 100 K in two consecutive NVT simulations (representing the canonical ensemble) of 5 ps each, imposing positional restraints of 100 kcal/mol Å^2^ on the protein-RNA complex. The temperature was further increased up to 200 K in a subsequent ∼100 ps MD run in the isothermal-isobaric ensemble (NPT), in which the restraint was reduced to 25 kcal/mol Å^2^. Finally, all restraints were released, and the systems were heated up to 300 K in a single NPT simulation of 500 ps. These simulations were performed using a 1 fs time step. The simulation time step was subsequently increased to 2 fs for further equilibration and production simulations. All bond lengths involving hydrogen atoms were constrained using the SHAKE algorithm. After ∼600 ps of equilibration, ∼10 ns of NPT simulation were carried out, allowing the systems’ density to stabilize around 1.01 g/cm^-3^. The temperature was kept constant at 300 K via Langevin dynamics (57), with a collision frequency γ = 1 ps^-1^. The pressure was controlled in the NPT simulations by coupling the system to a Berendsen barostat (58) at a reference pressure of 1 atm and with a relaxation time of 2 ps. Finally, each of the systems was simulated in the NVT ensemble in three replicates, reaching ∼1 μs for each replica, accumulating ∼12 μs of total sampling. All simulations were performed using the GPU-empowered version of the AMBER 20 simulation package (59). Analyses were performed over the aggregated multi-μs sampling collected for each of the studied complexes, offering a robust solid ensemble for the purposes of our analysis (detailed below and in **Supplementary Text 1**). Enhanced sampling simulations were also performed to compute the free energy profiles associated with the flipping of single nucleotide mismatches in the tgRNA (full details are reported in **Supplementary Text 1**).

### Dynamic network analysis and Signal-to-Noise Ratio

Graph-theory based network analysis was applied to characterize the allosteric pathways of communication (14). This analysis is well-suited for the characterization of allosteric mechanisms and for the identification of the most relevant communication routes between distal sites, as shown in a number of studies (60–63), including those performed by our research group (48–52). In dynamical networks, protein C⍺ atoms of protein and backbone P atoms of nucleotides, as well as N1 in purines, and N9 in pyrimidines, are represented as nodes, connected by edges weighted by the generalized correlations *GC_ij_* (details in **Supplementary Text 1**) according to:

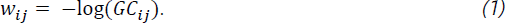

From the dynamical network, we estimated the efficiency of crosstalk between the crRNA spacer regions (i.e., nucleotides nt. 1-4, nt. 5-8, nt. 9-14, and nt. 15-18) and the catalytic residues (R472, H477, R1048, H1053) through a Signal-to-Noise Ratio (*SNR*) measure, introduced in our companion paper (Sinha et al. *under review*). *SNR* measures the preference of communication between predefined distant sites – i.e., the signal – over the remaining pathways in the network – i.e., the noise, estimating how allosteric pathways stand out (i.e., are favourable) over the entire communication network.

For the *SNR* calculation, we first computed the optimal (i.e., the shortest) and top five sub-optimal pathways (with longer lengths, ranked compared to the optimal path length) between all crRNA bases and the Cas13a residues, using well-established algorithms (details in the **Supplementary Text 1**). Then, the cumulative betweennesses of each pathway (*S_k_*) was calculated as the sum of the betweennesses of all the edges in that specific pathway:

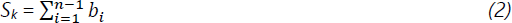

where *b_i_* is the edge betweenness (i.e., the number of shortest pathways that cross the edge, measuring the “traffic” passing through them) between node *i* and *i* + 1, and *n* is the number of edges in the *k*^*th*^ pathway. The distribution of *S_k_* between the crRNA bases and all protein residues was defined as the noise, whereas the distributions of *S_k_* between the crRNA nucleotide regions of interest (e.g., nt. 1-4) and the HEPN1-2 catalytic residues were considered as signals. Finally, the *SNR* corresponding to signals from each crRNA region to the HEPN1-2 catalytic core residues was computed as:

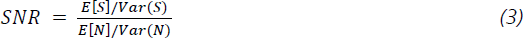

where *E(S)* and *Var(S)* correspond to the expectation and variance of the signal distribution respectively; and *E(N)/Var(N)* are the expectation/variance of the noise distribution. Notably, since SNR calculations are sensitive to the length of the communication pathways, the SNR was computed along shorter (edge count: 6-8), medium (edge count: 9-11), and longer (edge count: 12-14) path lengths (see **Supplementary Text 1**). The significance of the signal over the noise was estimated by computed using *Z*-score statistics with a two-tailed hypothesis. Finally, we also computed the ratio between the maximal *SNR* in the mismatched systems and in the system including perfectly matched (PM) crRNA: target-RNA duplex (*SNR_ratio_* = *SNR_max−mm_*/*SNR_max−pm_*). This comparison indicates whether single mismatches impact the strength of the communication with respect to the perfectly matched duplex. Full details on *SNR* calculations are reported in **Supplementary Text 1**. All networks were built using the Dynetan Python library (60). Path-based analyses were performed using NetworkX Python library (64).

### SARS-CoV-2 propagation

The following reagents were deposited by the Centers for Disease Control and Prevention and obtained through BEI Resources, NIAID, NIH: SARS-Related Coronavirus 2, isolate Hong Kong/VM20001061/2020, NR-52282; Isolate South Africa/KRISP-EC-K005321/2020 (B.1.351 lineage), NR-54008; isolate South Africa/KRISP-K005325/2020 (B.1.351 lineage), NR-54009; isolate USA/PHC658/2021 (Lineage B.1.617.2; Delta variant, NR-55611; and isolate USA/MD-HP20874/2021 (Lineage B.1.1.529; Omicron variant), NR-56461. SARS-CoV-2 was propagated (MOI of 0.1) and titered using 80% confluent African green monkey kidney epithelial Vero E6 cells (American Type Culture Collection, CRL-1586) or Vero-hACE2-TMPRSS2 cells (Vero AT) (BEI NR-54970) in Eagle’s Minimum Essential Medium (Lonza, 12-125Q) supplemented with 2% fetal bovine serum (FBS) (Atlanta Biologicals), 2 mM l-glutamine (Lonza, BE17-605E), and 1% penicillin (100 U/ml) and streptomycin (100 ug/ml) or puromycin (10 ug/ml) (Thermo Fisher, A11138-03). All isolates except Omicron were propagated and titered in Vero EG cells and using penicillin and streptomycin. The Omicron variant was propagated and titered in Vero AT cells using puromycin. Virus stock was stored at − 80°C. All work involving infectious SARS-CoV-2 was performed in the Biosafety Level 3 (BSL-3) core facility of the University of Rochester, with institutional biosafety committee (IBC) oversight.

### Tissue culture infectious dose assay, viral inactivation, and RNA extraction

Viral titers were determined using the tissue culture infectious dose (TCID) assay on triplicate wells of an 80% confluent monolayer of Vero E6 cells in a 96-well microtiter plate format using a 1:3 dilution factor; virus infection was assessed following 3-5 days of incubation at 37°C in a CO2 incubator by microscopic examination of cytopathic effects (CPE). The infectious dose (log10 TCID50/ml) was calculated using the Spearman-Kärber method (65,66). TCID50 of approximately seven logarithms per milliliter were used for RNA extractions. Infectious viral stocks were inactivated by a 1:3 dilution with TRI Reagent® (Zymo, R2050-1-200) immediately prior to RNA extraction. RNA was extracted using the Direct-zol RNA Miniprep Plus (Zymo, R2073) according to the manufacturer’s protocol, including on-column DNase 1 treatment.

### Viral and extracted sample preparation and RT-qPCR testing

To assess quality and relative quantity of viral RNA, RT-qPCR was performed using the Luna® SARS-CoV-2 RT-qPCR Multiplex Assay Kit (NEB) with the CDC-derived primers for N1 and N2 gene targets and the reaction was performed using the QuantStudio™ 5 System (ThermoFisher). Ct values for each viral strain used are reflected in **Table S6**.

For Cas13-cleavage assays, viral RNA was reversed transcribed using the High-Capacity cDNA Reverse Transcription Kit (Applied Biosystems). cDNA was amplified for the regions of interest using the primers listed in **Table S4**. The forward primers introduced a T7 RNAP promoter. PCR amplification was carried out using Q5® High-Fidelity DNA Polymerase (NEB) for at least 35 cycles, and annealing temperature of 60°C. Reaction products were visualized on a 2 % agarose gel with 0.05% (v/v) Ethidium Bromide and visualized with a Gel Doc XR+ imager (Bio-Rad Laboratories). PCR amplicons were column purified with the Monarch® PCR & DNA Cleanup Kit and eluted in 25 µL of Monarch® DNA Elution Buffer.

For the detection step, 1 µL of purified amplification product was added to 19 µL detection master mix (100 nM LbuCas13a:crRNA in cleavage buffer (20 mM HEPES-Na pH 6.8, 50 mM KCl, 5 mM MgCl_2_, 10 μg/mL BSA, 100 μg/mL tRNA, 0.01% Igepal CA-630 and 5% glycerol) with 1 U/µL murine RNase inhibitor (NEB), 0.1 µg/µL T7 RNA polymerase (purified in-house) and 1 mM of rNTP mix.

## RESULTS

### LbuCas13a exhibits differential sensitivity to mismatches in a position- and crRNA-length dependent manner

To study this phenomenon, we harnessed Cas13’s collateral-cleavage activity as a readout of Cas13 RNA-mediated HEPN-nuclease activation (15). Upon activation with a sufficiently complementary target RNA, Cas13 can non-specifically cleave quenched fluorescent reporters, resulting in increased fluorescent signal over time (**Figure 1A**). We first designed crRNAs targeting the same target-RNA with 16, 20 or 28 nucleotide (nt.) crRNA-spacer lengths (**Figure 1B**) and showed that Cas13a exhibits robust activation with these three crRNAs, albeit we observe a decrease in activity with a 16-nt. crRNA-spacer (**Figure 1C**).

**Figure 1.**
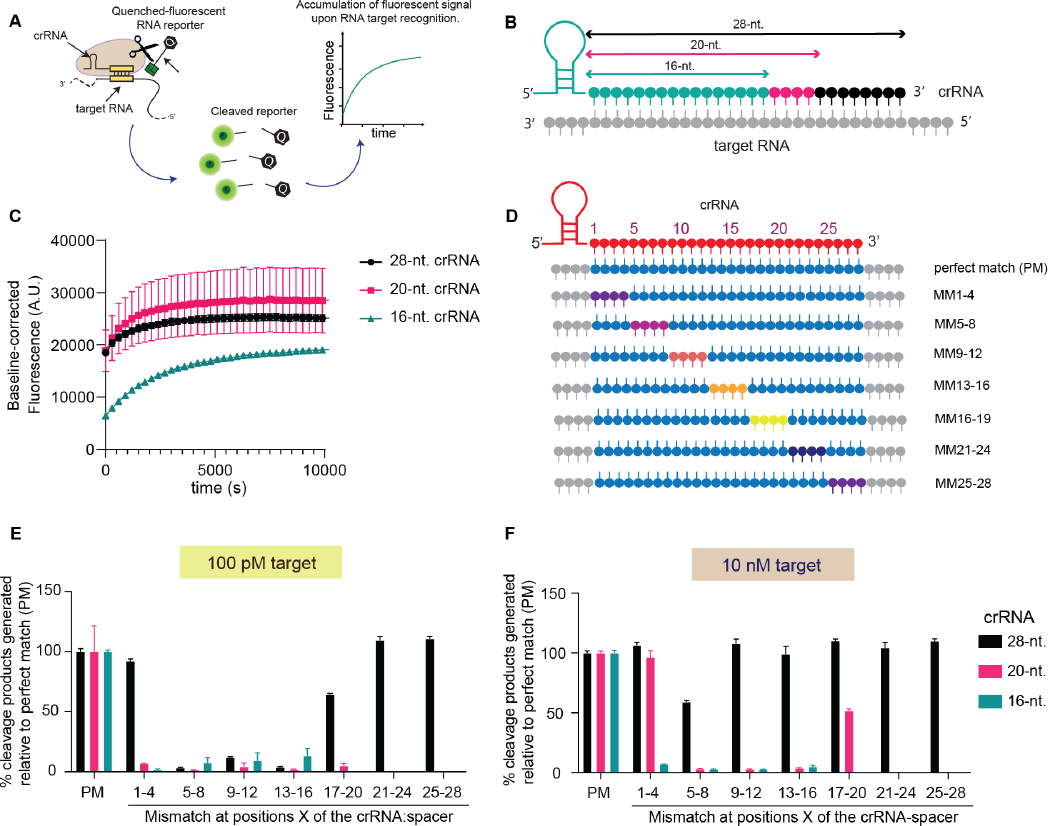
Cas13 exhibits differential sensitivity to mismatches in a position-dependent manner and it is modulated by crRNA spacer length. **A.** Schematic of a Cas13 RNA detection approach that harnesses the enzyme’s *trans*-ssRNA (collateral) cleavage activity for the cleavage of quenched fluorescent RNA reporters. **B.** Three different crRNA spacer lengths were used in the study of LbuCas13a nuclease activation: 16, 20 and 28 nucleotides. **C.** LbuCas13a reporter cleavage time-course with different spacer lengths against the same target RNA with 100 pM target concentration. **D.** Experimental design of the regions for which four consecutive mismatches (MM) were introduced between the crRNA and the target RNA. A perfect matched RNA is also used for reference (PM) **E.** LbuCas13a relative reporter cleavage efficiency after 1 hour using different crRNA spacer lengths and mismatched target-RNAs at 100 pM (4 consecutive mismatches) **F.** LbuCas13a relative reporter cleavage efficiency after 1 hour using different crRNA spacer lengths and mismatched target-RNAs at 10 nM (4 consecutive mismatches)

Given that LbuCas13a nuclease activity was maintained across all crRNA-spacer lengths tested, we then probed for sensitivity to mismatches at different regions of the crRNA-spacer:RNA-target region by introducing four consecutive mismatches across the complementarity region (**Figure 1D**) and assessed the amount of cleavage product generated after one hour incubation with different RNA target concentrations. While this experiment appears to be identical to previous experiments carried out by *Tambe et al.,* we noticed that this previous study incorporated an additional adenosine nucleotide at the 3’ end of the direct repeat (DR) in the crRNA, which inadvertently introduces an additional mismatch between the spacer and the target RNA (**Figure S1A-B),** and thus we decided to revisit these experiments using the canonical LbuCas13a DR (67) that does not include this additional adenosine. Of note, the mismatches in the target RNA were generated by replacing the nucleotide(s) in the target-RNA with the same nucleotide one present in the crRNA-spacer sequence, such that mismatch pair is made up of two of the same nucleotide. At 100 pM RNA target and targeting with a 28-nt. crRNA-spacer, ribonuclease activity could not be detected with mismatches between regions +5 to +13 relative the crRNA-spacer (**Figure 1E, S2A**). Interestingly, with shorter 16 or 20-nt. crRNA-spacer only a perfectly matched target-RNA was able to activate LbuCas13a (**Figure 1E, S2B-C**). We tested the same panel of mismatched target-RNAs using higher target-RNA concentration (10 nM) and observed that most of these cleavage defects were rescued at this high concentration when using a 28-nt. crRNA-spacer, with only a slightly decreased activity in region 5-8 (**Figure S2D**). The 20-nt. crRNA-spacer maintained RNase activity when there were mismatches in the 1-4 or 17-20 regions (**Figure 1F, S2E**), and the 16-nt. crRNA-spacer still required a perfect match for RNase activity (**Figure 1F, S2F**). However, as we saw in **Figure 1C** and **S2E,** using a crRNA with a 16-nt. spacer comes at a cost of total fluorescence magnitude and time to reach plateau, which could undermine sensitivity and assay times for diagnostic applications. As a result, we decided not to further pursue 16-nt. spacer design in this study.

Taken together, as expected, LbuCas13a displayed differential sensitivities to mismatches between the crRNA-spacer and its target-RNA in a position-dependent manner, and importantly, this sensitivity profile changes depending on the length of the crRNA-spacer used, with shorter crRNA-spacers being more sensitive to mismatches, and thus yielding higher specificity Cas13-crRNA complexes. It also cannot be overstated that higher concentrations of partially matched target RNA can in some cases still elicit nuclease activity, and this activity needs be considered when designing diagnostic assays, especially in context of pre-amplification of the RNA target, where fine control of the final concentration of target is difficult and can vary from sample to sample.

### Probing single nucleotide positions uncovers mismatch-sensitive hotspots in Cas13 that become more apparent with shorter crRNA-lengths

To further assess the effect of crRNA-spacer length on mismatch tolerance, we explored the effect that single nucleotide mismatches in the crRNA-target duplex have on Cas13 activation. We generated RNAs that contained a single nucleotide mismatch at each one of the first 20 nt. positions of the spacer (using the canonical LbuCas13a DR), performed trans-RNA cleavage assays and assessed the relative cleavage efficiencies compared to a perfectly complementary RNA target. For crRNA-spacer lengths of 28 and 20 nucleotides, single nucleotide mismatches across the length of the spacer did not impact Cas13 RNase activity at high target concentrations (10 nM) (**Figures 2A-B**).

**Figure 2.**
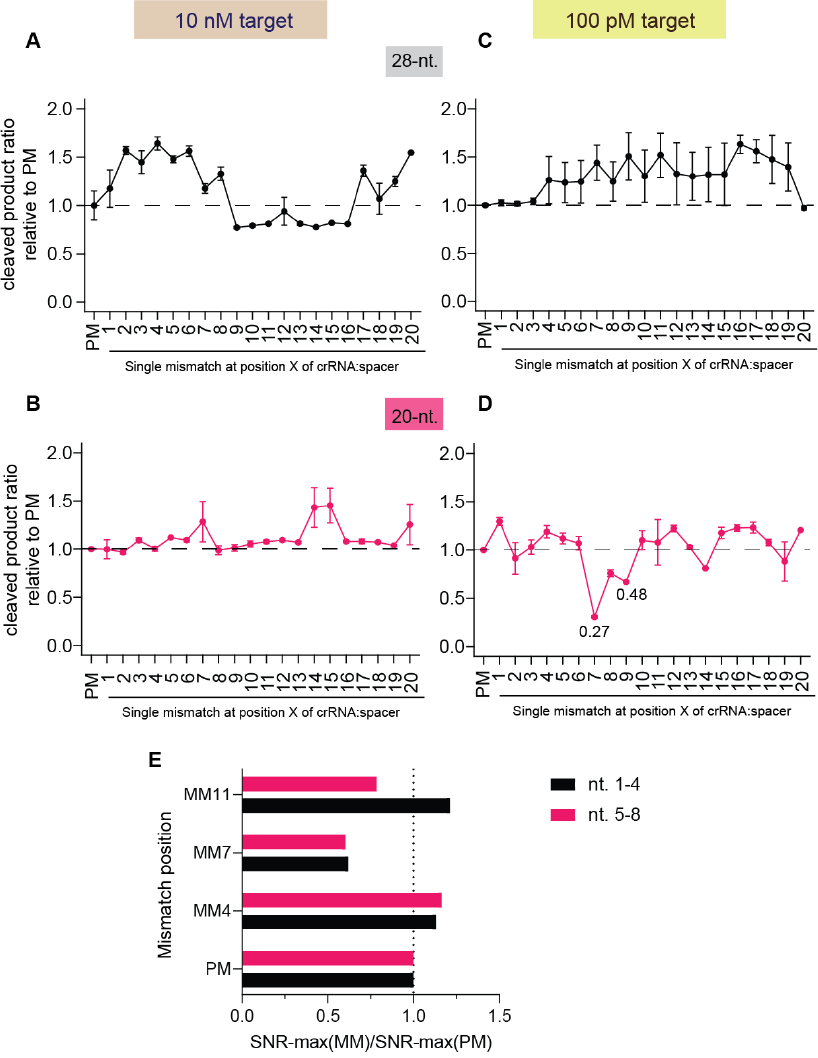
Probing single nucleotide positions uncovers mismatch-sensitive hotspots in Cas13. End-point (1 hour) LbuCas13a cleavage efficiencies of mismatched target-RNAs tiling at single nucleotide resolution across the target RNA compared to a perfect-matched (PM) ssRNA at two different target-RNA concentrations (10 nM and 100 pM) as follows: **A.** a 28 nucleotide spacer and 10 nM target; **B.** a 20 nucleotide spacer and 10 nM target; **C.** a 28 nucleotide spacer and 100 pM target; **D.** a 20 nucleotide spacer and 100 pM target; **E.** Ratio between the maximal Signal-to-Noise Ratio (*SNR*) in the Cas13a:crRNA:targer-RNA systems containing a PM target RNA and single mismatches at positions 4, 7 or 11; computed for spacer nt. 1-4 and nt. 5-8. Dashed lines indicate the ± 20% change with respect to the PM system.

When we lowered target-RNA concentrations to 100 pM, 28-nt. still yielded high Cas13a activity regardless of the presence of single-nucleotide mismatches in the target RNA (**Figure 2C**). On the other hand, when using a 20-nt. crRNA-spacer, single mismatches result in diminished RNase activation at several positions in the middle of the spacer (+7, +8, +9), and these mismatches resulted in up to a 70% reduction in activity, compared to a perfect-matched target-RNA under the same conditions (**Figure 2D**). It should be noted that compared to *Tambe et al.,* the location of mismatch-sensitive and tolerant regions was slightly different. In particular, this single mismatch profiling seems to indicate a subtle shift of one nucleotide in mismatch sensitivity, consistent with the different crRNA design (**Figure S3**).

Taken together, probing each base pair in the spacer:target-RNA duplex via mismatch analysis uncovers potentially useful mismatch sensitivities at specific regions of the spacer, however this is highly dependent on the length of the spacer as well as the target RNA concentration. In addition, 28-nt. spacers yield no single mismatch discriminatory capacity at the target-RNA concentrations tested.

### Investigating allosteric coupling associated with mismatch sensitivity using molecular dynamics simulations

We recently determined that certain regions of the crRNA-spacer: target-RNA duplex are most important for gating HEPN nuclease activation (45), and further explored this potential allosteric coupling using computational approaches (Sinha et al. *under review*). Our companion paper showed that target RNA binding acts as an allosteric activator of the HEPN nuclease domains and identified critical regions of LbuCas13b responsible for this information transfer (Sinha et al. *under review*). With this in mind, we wondered whether the differential, position-dependent mismatch sensitivity observed with 20-nt. length crRNA-spacers is also associated with perturbed allosteric coupling between the spacer nucleotides and catalytic residues of LbuCas13a, we conducted similar molecular dynamics (MD) simulations of the LbuCas13a complexes with single mismatches introduced at different locations within a 20-nt. spacer, as in our companion paper. MD simulations were carried out on four Cas13a: crRNA: target-RNA complexes containing either a perfectly matched crRNA: target-RNA duplex, or crRNA: target-RNA duplexes that contain a single mismatch at either spacer nucleotide position 4, 7, or 11. These positions were chosen either because they displayed a large loss of cleavage activity when mismatched (position +7) or no noticeable loss of cleavage activity (positions +4 and +11), as a negative control for our downstream analyses. Each system was simulated for ∼1 μs and in three replicates, collecting a multi-μs ensemble necessary for the analysis of the allosteric signaling (48–50).

We conducted graph-theory based network analysis to characterize the allosteric pathways of communication in the presence of single mismatches and compared these with the system with a perfectly matched crRNA-target RNA duplex. To estimate the communication efficiency between the crRNA spacer and the catalytic residues (R472, H477, R1048, H1053), we employed a Signal-to-Noise Ratio (SNR) measure, also used in our companion paper (Sinha et al. *under review*). The SNR measures the preference of communication between predefined distant sites – i.e., the signal – over the remaining pathways of comparable length in the network – i.e., the noise. The SNR thereby estimates how allosteric pathways stand out (i.e., are favourable) over the remaining noisy routes, with high *SNR* values indicating the preference for the network to communicate through the signal (see *Materials and Methods*).

To detect crRNA-spacer regions with preferred communication with catalytic residues, we performed SNR calculations considering signals sourcing from specific crRNA-spacer regions (i.e., nucleotides 1-4, 5-8, 9-14, 15-18 and 19-20) and sinking to the catalytic residues (R472, H477, R1048, H1053). The communication between all crRNA spacer nucleotides and all residues of the LbuCas13a protein was considered as the noise for all SNR calculations. As SNR calculation is sensitive to the pathlength of communication, which refers to the number of edges involved in the pathways connecting the spacer nucleotides with the catalytic residues, we evaluated SNR across different pathlengths. Specifically, we characterized the SNR for shorter (edge count: 6-8), medium (edge count: 9-11), and longer paths (edge count: 12-14) (see *Materials and Methods)* (**Figure S4**). The obtained SNR along any of the paths assesses the prevalence of the signal over the noise by determining the extent to which the signal distribution differs from the noise distribution. As shown in **Figure S4**, the perfectly matched crRNA-spacer system exhibits relatively higher SNR values for spacer 1-4-nt. along shorter and medium-length paths, as well as for 5-8-nt. along longer paths, compared to other regions.

To facilitate comparison, we compared the highest SNR observed in the no mismatch system with the highest SNR observed in the single mismatched systems, specifically corresponding to 1-4-nt. and 5-8-nt. across various pathlengths (**Figure 2E**). This comparison indicates whether the introduction of single mismatches has impacted the strength of communication between the spacer and the catalytic residues compared to the system without mismatch. The ratio between the highest SNRs in the single-mismatched and no-matched systems reveals that mismatches at positions 4 and 11 still result in a similar level of communication as the no-mismatch system, with perturbation within 20% of this system. In contrast, introducing a mismatch at position 7 results in a ∼40% reduction in SNR compared to no mismatches, both for 1-4-nt. And 5-8-nt. These results suggest that mismatches at position 7, which impact LbuCas13a nuclease cleavage rate, result in a loss of allosteric crosstalk between the spacer and catalytic residues, unlike the mismatches at 4 or 11 which do not result in a loss of cleavage activity experimentally nor a large loss in allosteric crosstalk in these simulations. Please see **Supplementary Text 1** and **Figure S6** for details on how we confirmed that the mismatched base conformations in our simulations reflect their expected energy minima.

### Structure-guided engineering of LbuCas13a gives rise to Cas13 variants with more stringent mismatch specificity profiles

Several different successful strategies to increase the sequence specificity of CRISPR-Cas enzymes have been employed in the past, including weakening the enzyme-target DNA interactions or slowing cleavage rates (68). The net effect is that the increasing difference between the dissociation and catalytic constant rates, allows the dissociation of non-perfect targets to be more favorable than cleavage, thus increasing discrimination. Applying this kinetic rationale, structure-guided engineering of CRISPR-Cas enzymes has yielded variants with high on-target cleavage and minimal off-target effects that improve their safety profile when used in research or therapeutic applications (69–72). Given our observations above and our recent computational study disclosing that the key R377, N378, and R973 residues gate the RNA-mediated allosteric HEPN activation (Sinha et al., *under review*), we hypothesized that by altering these allosteric communication pathways we could also alter the mismatch tolerance profile of LbuCas13a which, in turn, could facilitate the development of higher-fidelity Cas13 enzymes.

Specifically, we found that variants LbuCas13a^R377A^, LbuCas13a^N378A^, LbuCas13a^R973A^ have altered allostery communication pathways and we sought to explore the mismatch tolerance across the crRNA-spacer for each of these Cas13a variants. To this end, we overexpressed and purified these proteins as previously described (Methods) (**Figure S7A**) and performed cleavage assays either in the presence of a perfect matched target-RNA to our 28-nt. crRNA-spacer, or ssRNAs with four consecutive mismatches tiling across the crRNA-target RNA duplex (**Figure 3A**). End-point background-subtracted fluorescence measurements after one hour showed that each one of these Cas13a variants is more sensitive to mismatches compared to wild-type Cas13a (**Figure 3A**). For example, wild-type Cas13a still exhibited robust nuclease activation with mismatches at positions 1-4 or 17-20, where all Cas13a variants tested showed a significant reduction in nuclease activity with these mismatched target-RNAs. No significant change in apparent cleavage efficiency occurred in regions 21-24 and 25-28 when mismatches were present. These results suggest that the Cas13a variants have higher sensitivity to mismatches and thus may make suitable candidates for the development of higher-fidelity Cas13 enzymes for RNA-detection applications.

**Figure 3.**
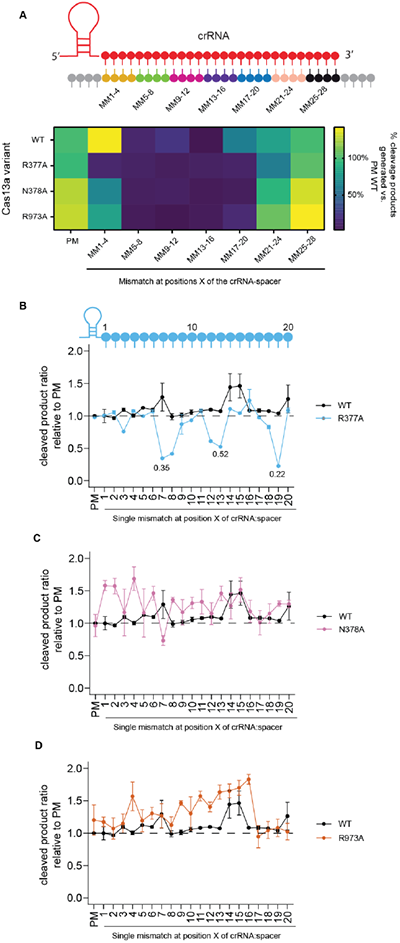
LbuCas13a variants show higher reporter assay specificity against mismatches between the crRNA-spacer and the target RNA. **A**. Heatmap of end-point fluorescence signal after 1 hour of LbuCas13a wild-type (WT) vs. variants using 10 nM of target RNA, either with no mismatches (PM) or 4 consecutive mismatches in the indicated regions. 28-nt. crRNA spacers were used. Results are background subtracted and normalized to values from WT LbuCas13a in the presence of PM RNA. **B-D.** Relative cleavage efficiencies for each LbuCas13a variant in the presence of a perfect match RNA (PM) or a single-nucleotide mismatched RNA (SM) at different positions relative to the crRNA and at 10 nM concentration and 20-nt. crRNA-spacer. Cleavage efficiency was normalized to wild-type (WT) LbuCas13a in the presence of PM RNA for each of the studied variants as follows: **B.** LbuCas13a^R377A^; **C.** LbuCas13a^N378A^; and **D.** LbuCas13a^R973A^.

Given, we saw no sensitivity to mismatches in regions 21-28 (**Figure 3A**) and our data in Figure 1E suggest that 20-nt. spacers are most appropriate with respect to generating new more specific Cas13 variants with single-nucleotide discrimination potential. To this end, we wanted to further understand the contributions of each single nucleotide in cleavage efficiency, and we used our previously generated RNAs with mismatches at each nucleotide position for every single position in the crRNA-target duplex. End-point normalized background-subtracted fluorescent values indicate that our LbuCas13a^N378A^ and LbuCas13a^R973A^ variants show higher sensitivity to single-nucleotide mismatches from 100 pM ssRNAs with 20 nucleotide spacers and display a more pronounced loss of activity profile across the crRNA-target compared to the wild-type LbuCas13a (**Figure S7B**).

Remarkably, LbuCas13a^R377A^ is not able to activate with the low 100 pM ssRNA concentration (**Figure S7B**) but at higher concentrations of ssRNA, activity with perfect-matched RNA is comparable to wild-type but single-pair mismatches at positions 7, 8, 12, 13, 19 within the crRNA-target RNA duplex particularly sensitive and is sufficient to result in loss of nuclease activity (**Figure 3B**). At this concentration, on the other hand, variants LbuCas13a^N378A^ and LbuCas13a^R973A^ do not exhibit any significant decrease in activity with single mismatches, and on the contrary, single mismatches at some positions seem to yield more robust activation (**Figure 3C-D**). We applied the same approach to 28-nt spacers and found that regardless of target concentration, 28-nt. crRNAs do not show sufficient discrimination of mismatched RNAs (**Figure S8A-B**).

Thus, altering residues that participate in the allosteric communication involved in HEPN nuclease activation gives rise to enzyme variants that display higher sensitivities to single nucleotide mismatches, although the degree of discriminatory power is position and spacer-length dependent. If using 20-nts. spacers, LbuCas13a^N378A^ and LbuCas13a^R973A^ variants are excellent at discriminating single-nucleotide mismatches at certain positions (mainly 7 and 19) when RNA target concentrations are low. If concentration of target is expected to be high, then LbuCas13a^R377A^ might make a strong candidate to distinguish single-nucleotide differences at those same positions.

### Deletion in the crRNA direct repeat contributes to the partial inhibition by anti-tag containing RNAs and result in better discrimination of mismatch containing RNAs

During our exploration in the literature of LbuCas13a crRNA architectures for different applications, it did not escape to our attention that the crRNA DR sequence used by *Meeske and Marraffini* for the study of ‘anti-tag’ RNA-mediated nuclease inhibition of Cas13a contains a deletion in the first adenine in the DR compared to other crRNA DR sequences commonly used for LbuCas13a (73) (**Figure 4A**). They reported that extended complementarity of the target RNA (forming an ‘anti-tag’) with the direct repeat of the crRNA of about 8 nucleotides results in inhibition of Cas13a activity despite having perfect complementarity in the spacer (**Figure 4B**). We wondered if this deletion in the DR might contribute at least partially to the observed inhibition of Cas13 cleavage in the presence of anti-tag containing RNAs. We changed the flanking regions of our target RNA with an anti-tag sequence for LbuCas13a crRNA (atgRNA) and used our original sequence without anti-tag (tgRNA). Additionally, we designed the same crRNAs as previously used but containing the A_-29_ deletion (del crRNA) and compared its activity relative to the full-length DR containing crRNA (WT crRNA). We then performed our trans-cleavage reporter assay in the presence of these crRNAs and targets (with and without anti-tag). The apparent cleavage efficiencies suggest that performing this truncation makes LbuCas13a more sensitive to inhibition by anti-tag containing RNAs, especially with low concentrations of target (100 pM) where the apparent cleavage activity is greatly reduced (**Figure 4C**). Higher concentrations of target RNA can overcome this inhibition.

**Figure 4.**
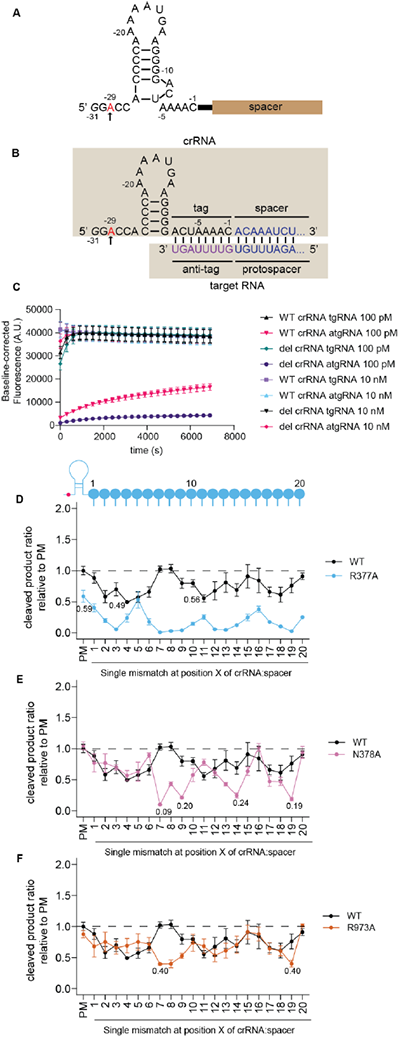
Deletion in the crRNA direct repeat contributes to the partial inhibition by anti-tag containing RNAs and result in better discrimination of mismatch-containing RNAs. **A.** Schematic of LbuCas13a crRNA sequence and structure. In red and pointed with an arrow, the adenine nucleotide that was deleted in Meeske and Marraffini (2018) and denoted as *del crRNA*. **B.** Schematic of LbuCas13a crRNA structure when pairing with an anti-tag containing RNA. The extended 5’ 8-nucleotide complementarity with the direct repeat results in partial inhibition of LbuCas13a nuclease activity. In red and pointed with an arrow, the adenine nucleotide that was deleted in Meeske and Marraffini (2018) and denoted as *del crRNA*. **C.** LbuCas13a reporter cleavage time-course with 20-nt. spacer of full-length (WT crRNA) or truncated crRNA (del crRNA) against the same target that contains an anti-tag (atgRNA) or not (tgRNA) with 100 pM or 10 nM final RNA target concentration as indicated. **D-F.** Relative cleavage efficiencies for each LbuCas13a variant in the presence of a perfect match RNA (PM) or a single-nucleotide mismatched RNA (SM) at different positions relative to the crRNA and at 10 nM concentration. The crRNA used contain the adenine deletion in the direct repeat and the spacer is 20 nucleotides long. Cleavage efficiency was normalized to wild-type (WT) LbuCas13a in the presence of PM RNA for each of the studied variants as follows: **D.** LbuCas13a^R377A^; **E.** LbuCas13a^N378A^; and **F.** LbuCas13a^R973A^.

Conversely, when using the crRNAs and sequences reported by *Meeske and Marraffini*, and restoring the DR to full length, the nuclease activity in the presence of high anti-tag RNA concentrations is rescued to the same levels as the RNA target (**Figure S9A-B**). Additionally, it appears that the deletion results in decreased nuclease activity even with targets that do not contain an anti-tag sequence.

While there might be sequence dependent effects mediating Cas13 activation, these assays suggest that truncating the direct repeat causes a subtle activation defect in Cas13a and makes it more sensitive to inhibition by anti-tag RNAs. Given the additional penalty imposed by this crRNA design, we hypothesized that using this truncated crRNA architecture (**Figure S10A**) may have an impact in the mismatch tolerance profile of LbuCas13a, increasing its mismatch discrimination ability. To investigate this, we performed additional cleavage assays with single-nucleotide mismatched target RNAs, for both wild-type LbuCas13a and the variants we investigated above, loaded with a truncated crRNA. For spacer lengths of 20-nt., a single nucleotide mismatch had greater impact on WT Cas13a activity at many positions across the crRNA-target region when the crRNA is truncated, even at high (10nM) target RNA concentrations (**Figure 4D-F**). At low (100 pM) target concentrations, the relative cleavage activity on WT Cas13a with mismatched target RNAs is very low, resulting in near complete loss of activity in most cases, particularly in the middle region of the spacer, for example, positions 7-9 and 14 (**Figure S10B**). Combining this truncated crRNA with a 20-nt. spacer and the variant LbuCas13a enzymes, the discrimination between a perfectly matched RNA and a mismatched one is further improved in some cases (**Figures 4D-F**). For example, LbuCas13a^R377A^ activity with a perfectly matched target-RNA (at 10 nM) is decreased about 50% compared to its wild-type counterpart (**Figure 4D**). Despite this apparent loss in activity, there is close to no activity of LbuCas13a^R377A^ in the presence of mismatches at most positions, particularly 3, 7-9, 12-14, 19 at high target concentrations (**Figure 4D**). For LbuCas13a^N378A^ and LbuCas13a^R973A^, robust activation with mismatched target-RNAs at high target-RNA concentrations is mostly observed, with the exception of a few positions, mainly positions 7 and 19 (**Figures 4E-F**). For LbuCas13a^N378A^, the presence of these mismatches in positions 7 or 19 resulted in almost no signal (**Figure 4E**), whereas for LbuCas13a^R973A^ only a partial loss of activation is observed (∼50%) (**Figure 4F**). Interestingly, at low concentrations of RNA target (100 pM), no nuclease activity is detected in any of the LbuCas13a variants even for a no mismatch RNA (**Figure S10B**), suggesting a decrease in sensitivity across all these variants when in combination with this truncated crRNA. Together, our data suggest that combining a 20-nt. truncated crRNA with LbuCas13a^N378A^ could be a promising candidate for the deployment of a much more specific Cas13-based diagnostic tool.

On the other hand, using 28-nt. spacers with a truncated DR with WT LbuCas13a does not result in single-nucleotide sensitivity, at either 100 pM or 10 nM target RNA (**Figure S10C-D**). If using our LbuCas13a variants, LbuCas13a^R377A^ is inactive at low target concentrations, even with a fully complementary RNA (**Figure S10C**), but at higher concentrations (10 nM), LbuCas13a^R377A^ is active and mismatches at positions 7 or 8 result in loss of cleavage activity that allows for single-nucleotide discrimination (**Figure S10D**). For LbuCas13a^N378A^ and LbuCas13a^R973A^, sensitivities at positions 7-10, 14 and 19 can be appreciated at 100 pM target RNA (**Figure S10C**). Raising the concentration of target to 10 nM abolishes this discriminatory ability with no significant sensitivity at any position (**Figure S10D**). If using 28-nt. crRNAs for diagnostic purposes is desired, combining the crRNA truncation with LbuCas13a^R377A^ would likely be the most sensible approach if the RNA concentrations are expected to be high.

Additionally, we used lateral flow readout to validate the potential for this discrimination approach to be adapted for point-of-care diagnostics. We compared the perfect-matched RNA and one with a mismatch at position 7 and compared the lateral flow readout using WT LbuCas13a and LbuCas13a^R377A^ (**Figure S11A**) with a full-length DR. Additionally using the truncated DR, we compared the same target RNAs with WT LbuCas13a, LbuCas13a^R377A^ and LbuCas13a^N378A^ (**Figure S11B**). In all cases, visual readout shows excellent discrimination using our new variants in all cases, unlike WT Cas13 that did not display sensitivity in the presence of mismatch 7 target RNA.

### SNP-detection of SARS-CoV-2 variants of concern uncovers additional crRNA design considerations

Given that together certain combinations of crRNA DR sequence variants and Cas13 protein variants are more sensitive to mismatches compared to WT LbuCas13a, we sought to test their performance in single nucleotide polymorphism (SNP) detection assays, which in turn could be deployed as a powerful point-of-care diagnostic test. This test could include infection diagnosis, genetic testing, detection of aberrant gene expression, cancer-related SNP or gene fusion detection, or epidemiological surveillance of pathogens. As a proof-of-concept, we designed assays against different regions of the Spike (S) protein transcript from SARS-CoV-2, for which mutations in this gene have resulted in the spread of new highly contagious and virulent SARS-CoV-2 strains. Particularly, we looked at the following amino acid mutational hotspots in the S transcripts that are signature mutations of some variants-of-concern (VOC): Beta (D80A), Delta (L452R) and Omicron (S477N + T478K) strains (**Figure 5A**). We designed crRNAs that were tailored to the ancestral strain or the VOC, such that mismatch(es) between the ancestral RNA and a VOC-specific crRNA (or vice versa) would result in discrimination by our Cas13a variants, (**Figure 5B**). We first tested these crRNAs by transcribing short RNAs with these regions of interest and performing cleavage measurements using a range of different mismatch combinations, Cas13 variants and crRNA DR designs based on the positive data above and arrived at several strategies that can be used in some cases to carry out more robust SNP discrimination than using WT LbuCas13a protein. (**Figures S12-13** see **Supplementary Text 2** for a detailed discussion of these data).

**Figure 5.**
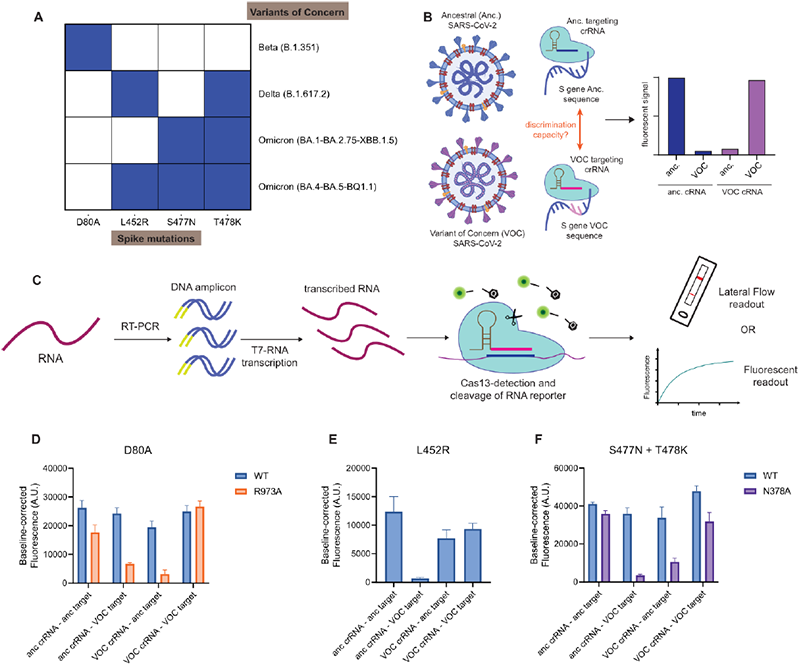
Combining highly specific Cas13a variants and rational crRNA design strategies can be deployed for SNP detection of SARS-CoV-2 strains. **A.** SARS-CoV-2 variants of concern and mutations relative to the ancestral strain assessed in this study. **B.** Schematic of Cas13a-based detection of SNP mutations in SARS-CoV-2 using a fluorescent readout with tailored crRNA designs; anc, ancestral; der, derived **C.** Schematic of a Cas13a RNA detection coupled to nucleic acid amplification. RNA is reverse transcribed into cDNA and T7 RNAP promoters are added during amplification. This amplified cDNA is used as template for T7 RNA polymerase transcription. The generated RNA products are detected by Cas13, and trans-cleavage of RNA reporters can be detected by either fluorescence or lateral flow, depending on the reporter design. **D.** Comparison of one-hour end-point fluorescence signal from WT and LbuCas13a^R973A^ when detecting a SARS-CoV-2 spike:D80S strain SNP (VOC target) or ancestral strain (anc. target) with a crRNA specific for the viral variant (VOC crRNA) or the ancestral virus (anc. crRNA). **E.** Comparison of one-hour end-point fluorescence signal from WT when detecting a SARS-CoV-2 spike:L452R strain SNP (VOC target) or ancestral strain (anc. target) with a crRNA specific for the viral variant (VOC crRNA) or the ancestral virus (anc. crRNA). **F.** Comparison of one-hour end-point fluorescence signal from WT and LbuCas13a^N378A^ when detecting a SARS-CoV-2 spike:S477N+T478K strain SNP (VOC target) or ancestral strain (anc. target) with a crRNA specific for the viral variant (VOC crRNA) or the ancestral virus (anc. crRNA).

We then harnessed these crRNA design strategies for the viral discrimination from cultured viral extracts. To do this, we employed a pre-amplification and detection strategy that couples amplification and T7 RNA polymerase transcription with our LbuCas13a cleavage assay and either a fluorescence or lateral flow read out to assay extracted RNA from SARS-CoV-2 virus generated via cell culture (**Figure 5C**). We found that for the detection of the D80A mutation using LbuCas13a^R973A^ and a crRNA (full DR) with a synthetic mismatch at position 19 and the discriminating mismatch at position 7 yielded strong recognition of the appropriate crRNA-target pairs but little activation in case the SNP of interest is present, unlike WT LbuCas13a that activates robustly in all cases (**Figure 5D, S14A-B, S15A**). For the L452R-causing SNP, we did not find a robust strategy as in these conditions the assay sensitivity seems to be low, but we observed that using WT LbuCas13a and crRNAs with the discriminating mismatch at position 19, there is discrimination for an ancestral-designed crRNA and a VOC target (**Figure 5E, S14C, S15B**). Finally, for Omicron variant detection (S477N + T478K) variant discrimination is achieved by using a crRNA where the discriminatory position is at nucleotide 19 for LbuCas13a^N378A^ compared to WT LbuCas13a (**Figure 5F, S14D-E, S15C**). Taken together, while our LbuCas13a SNP discrimination strategies are not completely generalizable yet, our data underscores that the addition of LbuCas13a variants and crRNA variants to the CRISPR diagnostics tool box offers an improvement in SNP discrimination ability compared to WT LbuCas13a and offers new opportunities for additional protein and crRNA engineering, and that sequence context likely plays a role in SNP discrimination power (see the next section for a further exploration of this idea) and thus multiple engineering strategies may be needed to enable easy to implement universal SNP discrimination.

### The type of mismatch and local sequence context modulates Cas13-mismatch tolerance

We noticed from our SARS-CoV-2-like discrimination assays using short RNA targets (**Supplementary Text 2** and **Figures S12-13**) that the degree of mismatch tolerance varies between the different RNA sequence targets tested in this study. Similar variability has been observed in other Cas13-based assays (32,40). Understanding whether there are additional crRNA or target RNA features that are contributing to these differences will allow researchers to make rational decisions about the most suitable crRNA design strategy to deploy for an RNA target of interest. We hypothesized that the base pairs participating in the mismatch might be a contributing factor, as well as nucleotide content in the crRNA-target duplex (e.g., G-C content).

To test the influence of interacting mismatched nucleotides, we choose the hyper-sensitive position 7 and obtained crRNAs and target-RNAs with the four possible nucleotides at that position and measured the relative cleavage efficiency, compared to a perfectly matched canonical base pair. Surprisingly, for either WT LbuCas13a at low target concentrations or LbuCas13a variants at high target concentrations, we observed that different mismatched pairs elicit different activation patterns (**Figure 6A, S16A-D**). For example, a C-C mismatch precludes Cas13 activation, but a G-G mismatch is very well tolerated, whereas G-A or C-U pairs have slightly different tolerances depending on their orientation in the crRNA-target.

**Figure 6.**
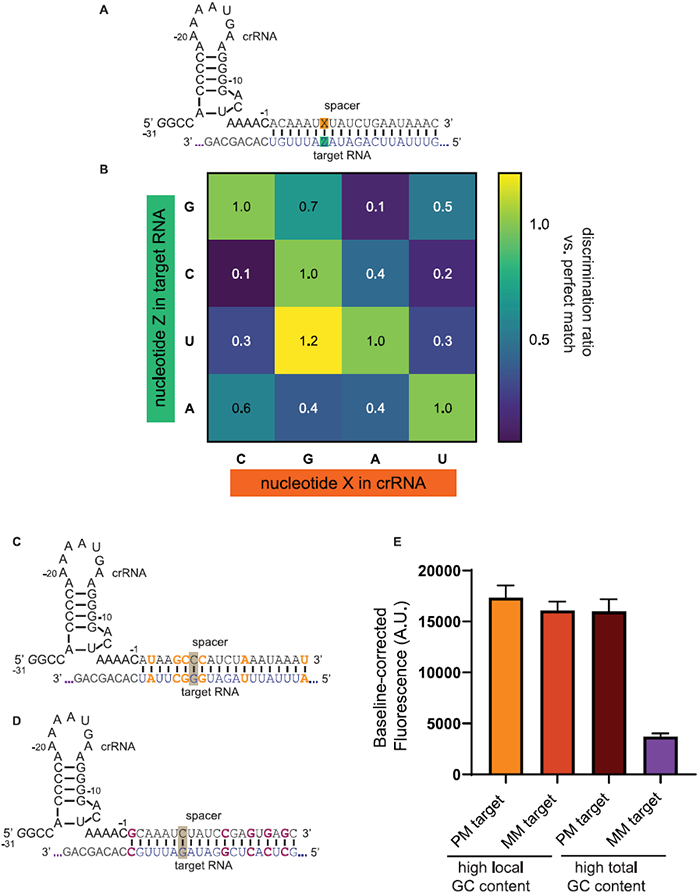
The type of mismatch and local sequence context modulates Cas13-mismatch tolerance. **A.** Schematic of crRNAs and target-RNAs used to study the specificity of LbuCas13a activation with all combinations of nucleotide base pairs at position 7 of the crRNA:target-RNA. **B.** Heatmap showing the ratio of cleaved products after one hour incubation with 100 pM of a target with a given nucleotide base pair combination at position 7 compared to its corresponding canonical base pair. **C.** Schematic of crRNA and target-RNA derived from our initial study by increasing the GC content around position 7 of the crRNA:target-RNA but compensating across the duplex to maintain the original 25% GC overall content. **D.** Schematic of crRNA and target-RNA derived from our initial study by increasing the total GC content from 25 to 50% but maintaining the original sequence context around position 7 of the crRNA:target-RNA. **E.** Comparison of one hour end-point fluorescence signal from LbuCas13a with 100 pM target, for the derived RNA sequences with different GC content, “high local GC content” corresponding to **C** and “high total GC content” corresponding to **D**. These measurements are performed with a perfectly-matched RNA target (PM) or containing a C-C mismatch at this position (MM).

Next, we tested whether the G-C content of the target RNA contributes to differences in mismatch tolerance. From our original target sequence, we derived two sequences: one with increased G-C content surrounding the sensitive position 7 but keeping the total original G-C content the same (25%) (**Figure 6B**), and other sequence where the bases around the 7^th^ position were maintained but the overall G-C content in the crRNA-target was raised from 25% to 50% (**Figure 6C**). Performing trans-cleavage assays in the presence of a perfect match and a mismatch at position 7 of the crRNA:target-RNA duplex, revealed that raising the GC composition around the mismatch resulted in robust nuclease activation. On the other hand, preserving the sequence context but raising the global GC content, mismatch sensitivity is still maintained (**Figure 6D**). Using higher concentrations of target RNA and our LbuCas13a variants yields similar differential tolerance based on sequence context and base pairs implicated in the mismatch (**Figure S17A-D**). Taken together, our mismatch and sequence-context cleavage analysis reveal that the nucleotide engaging in a mismatch and the local sequence context surrounding the mismatch modulates mismatch tolerance and thus, the cleavage specificity.

## DISCUSSION

The current COVID-19 pandemic has made apparent that scalability, affordability, flexibility, point-of-care (POC) availability, and fast-turnaround of diagnostic assays are key considerations and can influence policymaking, treatment options and public health strategies and alleviate morbidity (74,75). CRISPR-Cas systems – mainly Cas13 and Cas12 – have great potential to fulfill the need for a new generation of molecular diagnostic platforms. Efforts in recent years have focused on building these Cas13-based platforms for higher sensitivity, convenience, and multiplexing (16,17,31–34). In this study, we report a combination of strategies that can be used when designing Cas13a assays for SNP detection, as well as general considerations for any RNA-detection application with LbuCas13a. These include using shorter crRNA lengths, being aware of highly-sensitive (or insensitive) positions in the crRNA-target duplex, using specific LbuCas13a variants, employing crRNAs with modified direct repeats, accounting for the specific base-pair interactions or sequence contexts, and having an understanding that the concentration of target RNA can readily affect SNP detection performance.

Previous evidence has demonstrated that Cas13 shows differential activity and RNA-binding sensitivity to mismatches between the crRNA-target-RNA duplex in a position dependent manner (9,45). In particular, for LbuCas13a, four consecutive mismatches had different effects depending on their position in the spacer-target (45). By testing Cas13 nuclease activation with a canonical DR-spacer crRNA sequence and quadruple-mismatch target RNAs tiling across the crRNA-target, we were able to recapitulate the main conclusions of our previous study (45), although the location of mismatch-sensitive and tolerant regions was slightly different. In any instance, the middle region of the crRNA-target remains a hypersensitive region to tandem mismatches. The ability of Cas13 to occasionally engage in RNA cleavage with tandem mismatches, especially when those occur at the ends of the spacer-target complex and when target RNA concentrations are high, should be a central consideration when designing new RNA detection assays for Cas13a. This tolerance to mismatches is further exacerbated when using longer spacers (e.g., 28-nt.). While shorter crRNA-spacers display higher penalties to mismatches, it starts to come at a cost e.g., 16 nucleotide spacers show decreased cleavage rates relative to longer crRNA-spacers, and this can compromise the overall sensitivity of the assay. Taken all together, potential location of mismatches and length of the spacers should be first considerations for crRNA design and decisions should be made to tailor the crRNA design to the desired balance between sensitivity and specificity. Additionally, we performed single nucleotide mismatch profiling across the crRNA-target and measured the relative RNA cleavage activity compared to a non-mismatched target. Our profiling uncovered sensitive positions to mismatches, mainly positions 7 and 19. However, exploding these sensitivities is challenging, as high target RNA concentrations and/or longer spacers are able to overcome mismatch-induced cleavage defects. Similar observations pertaining to target concentrations have been made with Cas12a, where it has been observed that PCR cycles for target amplification needed to be controlled to prevent triggering off-target effects (29).

To harness and develop new ways of deploying LbuCas13a for SNP detection, we sought to obtain new enzyme variants with higher specificities. Our companion investigation (Sinha et al, under review) on structure-guided engineering of Cas13a proteins has yielded enzymes that are highly active but display more stringent mismatch tolerance profiles, thus making them more versatile and easier to deploy for RNA-detection applications. Thus, for SNP detection, our Cas13a variants are poised to gain higher levels of discrimination at the single-nucleotide level with more straightforward crRNA design principles. Specifically, these enzymes display sensitivities at specific critical crRNA-target-RNA regions that we can better exploit for improved SNP discrimination compared to WT Cas13a.

Relatedly, we show here that MD experiments coupled with a network analysis approach can be harnessed as a powerful tool to understand the effects of mismatches on the allosteric coupling between spacer: target-RNA binding and HEPN-nuclease activation. Our future work will continue to explore using these approaches to explore new and alternative molecular strategies to generate high fidelity Cas13 variants.

Furthermore, we have shown that subtle changes in the direct repeat of the crRNA make Cas13a more sensitive to inhibition by anti-tag containing RNAs or loss of activity with mismatched target RNAs. Specifically, by removing one adenine at the 5’ end of the crRNA, Cas13 shows strong activation but more likely to be inactivated by anti-tag containing RNAs, as previously reported (73). Additionally, we report that this truncation increases the mismatch tolerance for both WT LbuCas13a and our novel variants, thus making it an additional strategy for sequences that might be resistant to high specificity by other approaches. Our Future work will include a further exploration of crRNA design to determine whether there are additional modifications of the DR that allow for the generation of higher fidelity Cas13:crRNA complexes.

As proof-of-concept, we combined the principles we determined in this study for the discrimination of SNPs in SARS-CoV-2 VOCs and showed we can use them coupled to nucleic acid amplification for variant discrimination. Furthermore, we achieved better discrimination profiles compared to previously reported assays using LwaCas13a for these viral regions of interest (40). However, it did not escape our attention that there might be sequence-dependent effects that modulate the specificity threshold of Cas13a. We speculate that either the global sequence context or local sequence around the mismatch impacts the degree of the cleavage activity penalty from such mismatch, thus requiring the use of one or more synthetic mismatches or a combination of the strategies defined in this work if single-nucleotide specificity is needed. Moreover, we also noticed that the mismatched base pair type and the orientation (in the spacer vs. the target RNA) changed the extent of nuclease activation. Similar observations have been made where non-canonical base pairs can elicit robust Cas9 or Cas12 activation (29,76), and recent work in Cas13 demonstrated that G-U mismatches are the most tolerated (41,77). Future functional and structural studies of Cas13 enzymes will shed light on non-canonical base pairing tolerance, which, in turn, will further guide optimization efforts to select for base pairs within the target that can yield the highest discrimination power. Furthermore, we would also like to emphasize given that most Cas13-based RNA detection assays are amplification-based, special care should be given to make sure the working nucleic acid concentration obtained by amplification is above the limit of detection of the enzyme and assay, but also within ranges where the highest specificity may be achieved.

This study is limited by the use of a single ortholog, LbuCas13a. It is possible that position-specific mismatch sensitivities differ depending on the ortholog or that other Cas13 proteins naturally have higher specificity, and no further optimization and engineering is required. That said, our work provides a platform of guiding principles for diagnostic assay design for novel or known Cas13 variants that could be used for diagnostic purposes. While some of these strategies could lower the limit of detection of LbuCas13a, most Cas13-detection platforms require nucleic acid amplification, we are confident that a decrease in amplification-free limit of detection will have little detrimental effect in practice, if at all. Currently, there are no user-friendly crRNA design principles that guarantee SNP detection, and any potential design must be determined using a case-by-case basis, which can complicate the use of this technology for SNP diagnostics, especially when rapid customization is required (e.g., during outbreaks). The work presented here will aid in developing streamlined and accessible crRNA and Cas13 assay rules for various diagnostic applications.

## CONCLUSIONS

In sum, this study demonstrates an optimized Cas13a-based detection assay for detecting nucleotide variation in closely related sequences. We show that various crRNA design considerations are important for Cas13-detection development. We also present compelling evidence that the Cas13 variants we generated by studying the RNA-mediated allosteric activation of Cas13a are excellent candidates for highly specific detection tools, particularly for the detection of SNPs. Finally, we deployed all the lessons learned from this work for the detection of SARS-CoV-2 variants and showed their potential to fuel a new generation of Cas13a-based diagnostic tools.

## DATA AVAILABILITY

All relevant data are available in the manuscript and the supplementary materials. Data are also available on request from authors.

## SUPPLEMENTARY DATA

Supplementary Data are available at NAR online.

## AUTHOR CONTRIBUTIONS

**Adrian M. Molina Vargas**: Conceptualization, Data curation, Formal Analysis, Investigation, Methodology, Validation, Visualization, Writing-original draft. **Raven Osborn**: Resources, Data curation, Writing—review & editing. **Souvik Sinha**: Data curation, Formal Analysis, Writing—review & editing. **Pablo R. Arantes**: Data curation, Writing—review & editing. **Amun Patel**: Data curation, Writing—review & editing. **Stephen Dewhurst**: Resources, Writing—review & editing. **Giulia Palermo**: Supervision, Project administration, Funding acquisition, Writing—review & editing. **Mitchell R. O’Connell**: Conceptualization, Supervision, Project administration, Funding acquisition, Writing-original draft.

## ACKNOWLEDGEMENTS

The authors want to acknowledge the contributions of the UR CART/Center for Advanced Research Technologies Biosafety Level 3 (BSL3) facility, and the University of Rochester’s Institutional Biosafety Committee (IBC).

