## Supplemental Information and Figures for "New design strategies for ultra-specific CRISPR-Cas13a-based RNA-diagnostic tools with single-nucleotide mismatch sensitivity"

### Supplementary Material and Methods

#### Dynamic Network Analysis and Signal-to-Noise Ratio

Dynamic network analysis was performed to estimate the communication efficiency between the crRNA spacer regions and the catalytic cleft in the systems (1). In dynamical networks, protein residues are represented as nodes located at the $C_\alpha$ atoms, and nucleotides are represented by nodes located at the backbone P atoms, N1 atoms in purines, and N9 in pyrimidines (2). To determine whether nodes are involved in effective contacts, two criteria are used: ($i$) a distance cut-off of 4.5 Å between any two heavy atoms of two residues, and ($ii$) a frequency cut-off of 0.75, such that contacts are considered if formed for at least the 75 % of the simulation time. Nodes are connected by edges weighted by the generalized correlations $GC_{ij}$, based on Shannon's entropy estimation of mutual information (details on generalized correlations analysis are reported below) (3), according to:

$$w_{ij} = -\log(GC_{ij}) \quad (1)$$

From the dynamical network, we estimated the efficiency of crosstalk between the crRNA spacer regions (i.e., nucleotides nt. 1-4, nt. 5-8, nt. 9-14, and nt. 15-18) and the catalytic residues (R472, H477, R1048, H1053) through a Signal-to-Noise Ratio ($SNR$) measure, introduced in our companion paper (Sinha et al. *under review*). $SNR$ measures the preference of communication between predefined distant sites – i.e., the signal – over the remaining pathways in the network – i.e., the noise, estimating how allosteric pathways stand out (i.e., are favorable) over the entire communication network.

For the $SNR$ calculation, we first computed the optimal (i.e., the shortest) and top five sub-optimal pathways (with longer lengths, ranked compared to the optimal path length) between all crRNA bases and the Cas13a residues. Indeed, while the optimal path corresponds to the most likely mode of communication, suboptimal paths can also be crucial routes for communication transfer (1,4). Hence, in addition to the optimal path, we also considered the top five sub-optimal pathways for our $SNR$ analysis. Shortest paths calculations were performed using well-established algorithms. Specifically, the Floyd-Warshall algorithm was utilized to compute the optimal paths between the network nodes (5). The five sub-optimal paths were computed in rank from the shortest to the longest, using Yen's algorithm, which computes single-source $K$-shortest loop-less paths (i.e., without repeated nodes) for a graph with non-negative edge weights (6).

Then, the cumulative betweennesses of each pathway ($S_k$) was calculated as the sum of the betweennesses of all the edges in that specific pathway:

$$S_k = \sum_{i=1}^{n-1} b_i \qquad (2)$$

where $b_i$ is the edge betweenness (i.e., the number of shortest pathways that cross the edge, measuring the "traffic" passing through them) between node $i$ and $i+1$, and $n$ is the number of edges in the $k^{th}$ pathway. The distribution of $S_k$ between the crRNA bases and all protein residues was defined as the noise, whereas the distribution of $S_k$ between the crRNA nucleotide regions of interest (e.g., nt. 1-4) and the HEPN1-2 catalytic residues were considered as signals.

Since $S_k$ also depends on the number of edges present in the path, it can influence the $SNR$ measurement. Hence, to ensure consistency, we characterized the $SNR$ of communication pathways based on the number of edges present in the pathways: shorter (edge count: 6-8), medium (edge count: 9-11), and longer paths (edge count: 12-14). Toward this aim, we determined the stratification of path lengths through the kernel density estimation of the number of edges in the optimal and top five sub-optimal pathways that communicate the RNA spacer regions, nt. 1-4, nt. 5-8, nt. 9-14, nt. 15-18 with the catalytic residues in the system containing a perfectly matched crRNA: target-RNA duplex (**Figure S5**).

Finally, the $SNR$ corresponding to signals from each crRNA region to the HEPN1-2 catalytic core residues was computed as:

$$SNR = \frac{E[S]/Var(S)}{E[N]/Var(N)} \qquad (3)$$

where $E(S)$ and $Var(S)$ correspond to the expectation and variance of the signal distribution respectively; and $E(N)/Var(N)$ are the expectation/variance of the noise distribution.

To provide the significance of the signal over the noise, we used a general approach based on $p$-value calculation. Our goal was to test the hypothesis that the signal is an outlier of the noise distribution. We can construct a best-fit probability distribution based on the noise data by treating our variable (the sum of betweennesses) as stochastic. Treating the signal as a sample from this population, we can assess the rarity of that sample's mean when randomly collecting samples of the same size. This rarity is defined as the $p$-value of the sample. The $p$-value is then computed using the $Z$-score:

$$Z_{signal} = \frac{E[S]-E[N]}{\sigma[N]/\sqrt{n}} \tag{4}$$

where $\sigma[N]$ corresponds to the standard deviation of the noise, and $n$ is the number of signals. Here, we found that the best-fit distributions for the noise follow a log-normal distribution. Hence, we took the logarithm of the signals and noise to transform the data points into a normal distribution. From this transformed distribution, we obtained the mean and standard deviations for the calculation of the $Z_{signal}$. All networks were built using the Dynetan Python library (1). Path-based analyses were performed using NetworkX Python library (7) .

**Generalized correlation analysis**

Generalized Correlation ($GC$) analysis was used to describe the correlations between pairs of atoms and to construct dynamical networks. With this analysis, the correlation between atom pairs is described independently on the relative orientation of their respective fluctuations, capturing non-linear contributions to correlations. The generalized correlation coefficients are derived from mutual information, $MI$, a measure of independence between random variables, estimated for pair of nodes using their positional record in the simulated trajectory (3) .

In $GC$ analysis, two variables ($x_i$, $x_j$) can be considered correlated when their joint probability distribution, $p(x_i, x_j)$, is smaller than the product of their marginal distributions, $p(x_i) \cdot p(x_j)$. The $MI$ between $x_i$ and $x_j$ is defined as function of $p(x_i, x_j)$ and $p(x_i) \cdot p(x_j)$ according to:

$$MI\left[x_i, x_j\right] = \iint p(x_i, x_j)\, ln \frac{p(x_i, x_j)}{p(x_i) \cdot p(x_j)} dx_i dx_j \tag{5}$$

The $MI$ is closely related to the definition of the Shannon entropy, $H[x]$, i.e., the expectation value of a random variable $x$, having a probability distribution $p(x_i)$

$$H[x] = -\int p(x)\, ln\, p(x) dx \tag{6}$$

and it can be computed as:

$$MI\,[x_i,\, x_j] = H\,[x_i] + H\,[x_j] - H\,[x_i, x_j] \tag{7}$$

where $H\,[x_i]$ and $H\,[x_j]$ are the marginal Shannon entropies, and $H\,[x_i, x_j]$ is the joint entropy. Since $MI$ varies from 0 to +∞, normalized generalized correlation ($GC$) coefficients, ranging from 0 (independent variables) to 1 (fully correlated variables), are defined as:

$$GC\,[x_i,\, x_j] = \left\{1 - e^{-\frac{2MI[x_i, x_j]}{d}}\right\}^{-\frac{1}{2}} \tag{8}$$

where $d = 3$ is the dimensionality of $x_i$ and $x_j$. In the present work, $GCs$ were computed using the recent high-performance $GC$-based dynamical network analysis tool by Luthey-Schulten (2). $GC$ analysis has been performed by considering the $C_\alpha$ atoms for the protein residues and for the nucleotides the backbone P atoms, N1 atoms in purines, and N9 in pyrimidines (2).

**Enhanced sampling simulations**

To compute the free energy profiles associated with the flipping of single mismatches in the tgRNA, we performed enhanced sampling simulations, using the Umbrella Sampling (US) method (8). The reaction coordinate (RC) to describe base flipping was selected as the pseudo dihedral angle θ formed by the centres of mass (COM) of four groups of atoms: ($a$) the heavy atoms of the 5′ base-pair adjacent to the flipping base, ($b$) the sugar moiety of the 5′ base, ($c$) the sugar moiety of the flipping base, and ($d$) the flipping base (**Figure S6A**). This RC describes the flipping of the mismatched base relative to the preceding matched base pair, and was previously used in computational studies of base flipping (9,10). The RC ranges from −180° to +180.0° and was sampled in 10° steps (i.e., 36 US windows). Each window was sampled for ~15 ns reaching convergence of the free energy profiles (**Figure S6D-F**), and accumulating ~540 ns of sampling for each system. US simulations were initiated starting from the structures equilibrated after

500 ps of MD in the NPT ensemble, in which the configuration of the base-pairs at various mismatched locations was defined by $\theta$ ~20.0° - 30.0° (~0.35 – 0.5 rad). For each window, the starting structures were generated by carrying out 40 ps-long US runs with a force constant of 2000 kcal/mol per rad$^2$. The resulting structure from each simulation was then utilized as the starting structure for the subsequent window in the sampling process. For production runs, the harmonic force constant was set to 200 kcal/mol per rad$^2$. The free energy profiles for the flipping process were subsequently derived using the Weighted Histogram Analysis Method (WHAM)(11). WHAM analysis was carried out using a window width of 2°, finer than the US window width of 10°, with a convergence criterion of 0.001 kcal/mol. Uncertainties of the obtained free energy profiles were obtained from the profiles calculated at 1 ns interval, extracted from the final 5 ns of 15 ns window, at each grid point of the Dihedral RC. All WHAM calculations were performed using the Grossfield code [Grossfield, Alan, "WHAM: the weighted histogram analysis method", version 2.0.11, http://membrane.urmc.rochester.edu/wordpress/?page_id=126](12).

**Table S1. Oligonucleotides used for site-directed mutagenesis.**

| name | sequence | use |
|---|---|---|
| AM78_Lbu_R377A_ATH_fwd | AAGCGTTTCTTGCCAACATCATTGGGGT | site-directed mutagenesis |
| AM79_Lbu_N378A_ATH_fwd | AAGCGTTTCTTCGCGCCATCATTGGGGT | site-directed mutagenesis |
| AM80_Lbu_377_378_ATH_rev | CATTCTGACGGTTGCGGGCGAT | site-directed mutagenesis - rev for both AM78 and AM79 |
| AM89_Lbu_R973A_ATH_rev | GAAGCGCAGATCGGCTTCCCAAATTG | site-directed mutagenesis |
| AM90_Lbu_R973A_ATH_fwd | CGCCTTAAAGGTGAGTTCCCAGAAAACCAAT | site-directed mutagenesis |
| AM85_Lbu_973_seq_fwd | AGGACTATGAAAGTTACAAGCAAGCT | Sanger sequencing primer |
| AM86_Lbu_377_378_seq_fwd | TTGACACGTACGTCCGTAATTGT | Sanger sequencing primer |

**Table S2. Oligonucleotides used for *in-vitro* RNA transcription.**

| name | sequence |
|---|---|
| MOC-626_T7_opt | GGCGTAATACGACTCACTATAGG |
| MOC-1226_longer_AM_Liu_Lbu_target_IVTtemp | GTGTGGGCTTCTGCTGTGACAAATCTATCTGAATAAACTCTTCTTCTTGGTTTCCCtatagtgagtcgtattacgcc |
| MOC-1227_longer_AM_Liu_Lbu_antitag_IVTtemp | GTGTGGGCTTACTAAAACACAAATCTATCTGAATAAACTCTTCTTCTTGGTTTCCCtatagtgagtcgtattacgcc |
| AM91_Lbu_Liu_target_MM25:28 | GTGTGGGCTTCTGCTGTGG ACAAATCTATCTGAATAAACTCTTGAAG TTGGTTTCCCtatagtgagtcgtattacgcc |
| AM92_Lbu_Liu_target_MM21:24 | GTGTGGGCTTCTGCTGTGG ACAAATCTATCTGAATAAACAGAACTTC TTGGTTTCCCtatagtgagtcgtattacgcc |
| AM93_Lbu_Liu_target_MM17:20 | GTGTGGGCTTCTGCTGTGG ACAAATCTATCTGAATTTTGTCTTCTTC TTGGTTTCCCtatagtgagtcgtattacgcc |
| AM94_Lbu_Liu_target_MM13:16 | GTGTGGGCTTCTGCTGTGG ACAAATCTATCTCTTAAAACTCTTCTTC TTGGTTTCCCtatagtgagtcgtattacgcc |
| AM95_Lbu_Liu_target_MM9:12 | GTGTGGGCTTCTGCTGTGG ACAAATCTTAGAGAATAAACTCTTCTTC TTGGTTTCCCtatagtgagtcgtattacgcc |
| AM96_Lbu_Liu_target_MM5:8 | GTGTGGGCTTCTGCTGTGG ACAATAGAATCTGAATAAACTCTTCTTC TTGGTTTCCCtatagtgagtcgtattacgcc |
| AM97_Lbu_Liu_target_MM1:4 | GTGTGGGCTTCTGCTGTGG TGTTATCTATCTGAATAAACTCTTCTTC TTGGTTTCCCtatagtgagtcgtattacgcc |
| AM108_Lbu_Liu_target_SM1 | GTGTGGGCTTCTGCTGTGG tCAAATCTATCTGAATAAACTCTTCTTC TTGGTTTCCCtatagtgagtcgtattacgcc |
| AM109_Lbu_Liu_target_SM2 | GTGTGGGCTTCTGCTGTGG AgAAATCTATCTGAATAAACTCTTCTTC TTGGTTTCCCtatagtgagtcgtattacgcc |
| AM110_Lbu_Liu_target_SM3 | GTGTGGGCTTCTGCTGTGG ACtAATCTATCTGAATAAACTCTTCTTC TTGGTTTCCCtatagtgagtcgtattacgcc |
| AM111_Lbu_Liu_target_SM4 | GTGTGGGCTTCTGCTGTGG ACAtATCTATCTGAATAAACTCTTCTTC TTGGTTTCCCtatagtgagtcgtattacgcc |

| | |
|---|---|
| AM112_Lbu_Liu_target_SM5 | GTGTGGGCTTCTGCTGTGG ACAAtTCTATCTGAATAAACTCTTCTTC TTGGTTTCCCtatagtgagtcgtattacgcc |
| AM113_Lbu_Liu_target_SM6 | GTGTGGGCTTCTGCTGTGG ACAAAaCTATCTGAATAAACTCTTCTTC TTGGTTTCCCtatagtgagtcgtattacgcc |
| AM114_Lbu_Liu_target_SM7 | GTGTGGGCTTCTGCTGTGG ACAAATgTATCTGAATAAACTCTTCTTC TTGGTTTCCCtatagtgagtcgtattacgcc |
| AM115_Lbu_Liu_target_SM8 | GTGTGGGCTTCTGCTGTGG ACAAATCaATCTGAATAAACTCTTCTTC TTGGTTTCCCtatagtgagtcgtattacgcc |
| AM116_Lbu_Liu_target_SM9 | GTGTGGGCTTCTGCTGTGG ACAAATCTtTCTGAATAAACTCTTCTTC TTGGTTTCCCtatagtgagtcgtattacgcc |
| AM117_Lbu_Liu_target_SM10 | GTGTGGGCTTCTGCTGTGG ACAAATCTAaCTGAATAAACTCTTCTTC TTGGTTTCCCtatagtgagtcgtattacgcc |
| AM118_Lbu_Liu_target_SM11 | GTGTGGGCTTCTGCTGTGG ACAAATCTATgTGAATAAACTCTTCTTC TTGGTTTCCCtatagtgagtcgtattacgcc |
| AM119_Lbu_Liu_target_SM12 | GTGTGGGCTTCTGCTGTGG ACAAATCTATCaGAATAAACTCTTCTTC TTGGTTTCCCtatagtgagtcgtattacgcc |
| AM120_Lbu_Liu_target_SM13 | GTGTGGGCTTCTGCTGTGG ACAAATCTATCTcAATAAACTCTTCTTC TTGGTTTCCCtatagtgagtcgtattacgcc |
| AM121_Lbu_Liu_target_SM14 | GTGTGGGCTTCTGCTGTGG ACAAATCTATCTGtATAAACTCTTCTTC TTGGTTTCCCtatagtgagtcgtattacgcc |
| AM122_Lbu_Liu_target_SM15 | GTGTGGGCTTCTGCTGTGG ACAAATCTATCTGAtTAAACTCTTCTTC TTGGTTTCCCtatagtgagtcgtattacgcc |
| AM123_Lbu_Liu_target_SM16 | GTGTGGGCTTCTGCTGTGG ACAAATCTATCTGAAaAAACTCTTCTTC TTGGTTTCCCtatagtgagtcgtattacgcc |
| AM124_Lbu_Liu_target_SM17 | GTGTGGGCTTCTGCTGTGG ACAAATCTATCTGAATtAACTCTTCTTC TTGGTTTCCCtatagtgagtcgtattacgcc |
| AM125_Lbu_Liu_target_SM18 | GTGTGGGCTTCTGCTGTGG ACAAATCTATCTGAATAtACTCTTCTTC TTGGTTTCCCtatagtgagtcgtattacgcc |
| AM126_Lbu_Liu_target_SM19 | GTGTGGGCTTCTGCTGTGG ACAAATCTATCTGAATAAtCTCTTCTTC TTGGTTTCCCtatagtgagtcgtattacgcc |
| AM127_Lbu_Liu_target_SM20 | GTGTGGGCTTCTGCTGTGG ACAAATCTATCTGAATAAAgTCTTCTTC TTGGTTTCCCtatagtgagtcgtattacgcc |

| | |
|---|---|
| AM160_SARS_COV2_S_D80_WT | CATTAAATGGTAGGACAGGGTTATCAAACCTCTTAGTACCATTGGTCCCAGAGACCCtatagtgagtcgtattacgcc |
| AM161_SARS_COV2_S_D80_BETA | CATTAAATGGTAGGACAGGGTTAGCAAACCTCTTAGTACCATTGGTCCCAGAGACCCtatagtgagtcgtattacgcc |
| AM164_SARS_COV2_S_L452_WT | TTAGACTTCCTAAACAATCTATACAGGTAATTATAATTACCACCAACCTTAGAATCCtatagtgagtcgtattacgcc |
| AM165_SARS_COV2_S_L452_DELTA | TTAGACTTCCTAAACAATCTATACCGGTAATTATAATTACCACCAACCTTAGAATCCtatagtgagtcgtattacgcc |
| AM168_SARS_COV2_S_S477_WT | AAACCTTCAACACCATTACAAGGTGTGCTACCGGCCTGATAGATTTCAGTTGAAACCtatagtgagtcgtattacgcc |
| AM169_SARS_COV2_S_S477_OMICRON | AAACCTTCAACACCATTACAAGGTTTGTTACCGGCCTGATAGATTTCAGTTGAAACCtatagtgagtcgtattacgcc |

### Table S3. RNAs used in this study.

| Name / Purpose | sequence 5' - 3' | type |
|---|---|---|
| MOC-1264_Lbu_mat_crRNA_Liu_28nt | GGACCACCCCAAAAAUGAAGGGGACUAAAACACAAAUCUAUCUGAAU AAACUCUUCUUC | crRNA |
| MOC-1265_Lbu_mat_crRNA_Liu_20nt | GGACCACCCCAAAAAUGAAGGGGACUAAAACACAAAUCUAUCUGAAU AAAC | crRNA |
| MOC-1266_Lbu_mat_crRNA_Liu_16nt | GGACCACCCCAAAAAUGAAGGGGACUAAAACACAAAUCUAUCUGAAU | crRNA |
| AM208_Liu_mut_DR_28 | GGCCACCCCAAAAAUGAAGGGGACUAAAACACAAAUCUAUCUGAAUA AACUCUUCUUC | crRNA |
| AM211_Liu_mut_DR_Lbu_20 | GGCCACCCCAAAAAUGAAGGGGACUAAAACACAAAUCUAUCUGAAUA AAC | crRNA |
| MOC-1284_Liu_long_target_IDT_RNA | GGGAAACCAAGAAGAAGAGTTTATTCAGATAGATTTGTCACAGCAGAAG CCCACAC | target |
| MOC-1285_Liu_long_anti-target_IDT_RNA | GGGAAACCAAGAAGAAGAGTTTATTCAGATAGATTTGTGTTTTAGTAAGC CCACAC | target |
| Liu_Perfect_match_target_RNA | GGGAAACCAA GAAGAAGAGUUUAUUCAGAUAGAUUUGU CACAGCAGAAGCCCACAC | target |
| Liu_target_RNA_MM1-4 | GGGAAACCAA GAAGAAGAGUUUAUUCAGAUAGAUAACA CACAGCAGAAGCCCACAC | target |
| Liu_target_RNA_MM5-8 | GGGAAACCAA GAAGAAGAGUUUAUUCAGAUUCUAUUGU CACAGCAGAAGCCCACAC | target |
| Liu_target_RNA_MM9-12 | GGGAAACCAA GAAGAAGAGUUUAUUCUCUAAGAUUUGU CACAGCAGAAGCCCACAC | target |
| Liu_target_RNA_MM13-16 | GGGAAACCAA GAAGAAGAGUUUUAAGAGAUAGAUUUGU CACAGCAGAAGCCCACAC | target |
| Liu_target_RNA_MM17-20 | GGGAAACCAA GAAGAAGACAAAAUUCAGAUAGAUUUGU CACAGCAGAAGCCCACAC | target |
| Liu_target_RNA_MM21-24 | GGGAAACCAA GAAGUUCUGUUUAUUCAGAUAGAUUUGU CACAGCAGAAGCCCACAC | target |
| Liu_target_RNA_MM25-28 | GGGAAACCAA CUUCAAGAGUUUAUUCAGAUAGAUUUGU CACAGCAGAAGCCCACAC | target |

| | | |
|---|---|---|
| Liu_target_RNA_SM1 | GGGAAACCAA GAAGAAGAGUUUAUUCAGAUAGAUUUGa CACAGCAGAAGCCCACAC | target |
| Liu_target_RNA_SM2 | GGGAAACCAA GAAGAAGAGUUUAUUCAGAUAGAUUUcU CACAGCAGAAGCCCACAC | target |
| Liu_target_RNA_SM3 | GGGAAACCAA GAAGAAGAGUUUAUUCAGAUAGAUUaGU CACAGCAGAAGCCCACAC | target |
| Liu_target_RNA_SM4 | GGGAAACCAA GAAGAAGAGUUUAUUCAGAUAGAUaUGU CACAGCAGAAGCCCACAC | target |
| Liu_target_RNA_SM5 | GGGAAACCAA GAAGAAGAGUUUAUUCAGAUAGAaUUGU CACAGCAGAAGCCCACAC | target |
| Liu_target_RNA_SM6 | GGGAAACCAA GAAGAAGAGUUUAUUCAGAUAGUUUUGU CACAGCAGAAGCCCACAC | target |
| Liu_target_RNA_SM7 | GGGAAACCAA GAAGAAGAGUUUAUUCAGAUAcAUUUGU CACAGCAGAAGCCCACAC | target |
| Liu_target_RNA_SM8 | GGGAAACCAA GAAGAAGAGUUUAUUCAGAUUGAUUUGU CACAGCAGAAGCCCACAC | target |
| Liu_target_RNA_SM9 | GGGAAACCAA GAAGAAGAGUUUAUUCAGAaAGAUUUGU CACAGCAGAAGCCCACAC | target |
| Liu_target_RNA_SM10 | GGGAAACCAA GAAGAAGAGUUUAUUCAGUUAGAUUUGU CACAGCAGAAGCCCACAC | target |
| Liu_target_RNA_SM11 | GGGAAACCAA GAAGAAGAGUUUAUUCAcAUAGAUUUGU CACAGCAGAAGCCCACAC | target |
| Liu_target_RNA_SM12 | GGGAAACCAA GAAGAAGAGUUUAUUCUGAUAGAUUUGU CACAGCAGAAGCCCACAC | target |
| Liu_target_RNA_SM13 | GGGAAACCAA GAAGAAGAGUUUAUUgAGAUAGAUUUGU CACAGCAGAAGCCCACAC | target |
| Liu_target_RNA_SM14 | GGGAAACCAA GAAGAAGAGUUUAUaCAGAUAGAUUUGU CACAGCAGAAGCCCACAC | target |
| Liu_target_RNA_SM15 | GGGAAACCAA GAAGAAGAGUUUAaUCAGAUAGAUUUGU CACAGCAGAAGCCCACAC | target |
| Liu_target_RNA_SM16 | GGGAAACCAA GAAGAAGAGUUUUUUCAGAUAGAUUUGU CACAGCAGAAGCCCACAC | target |

| | | |
|---|---|---|
| Liu_target_RNA_SM17 | GGGAAACCAA GAAGAAGAGUUaAUUCAGAUAGAUUUGU CACAGCAGAAGCCCACAC | target |
| Liu_target_RNA_SM18 | GGGAAACCAA GAAGAAGAGUaUAUUCAGAUAGAUUUGU CACAGCAGAAGCCCACAC | target |
| Liu_target_RNA_SM19 | GGGAAACCAA GAAGAAGAGaUUAUUCAGAUAGAUUUGU CACAGCAGAAGCCCACAC | target |
| Liu_target_RNA_SM20 | GGGAAACCAA GAAGAAGAcUUUAUUCAGAUAGAUUUGU CACAGCAGAAGCCCACAC | target |
| AM160_SARS_COV2_S_D80_WT | GGGTCTCTGGGACCAATGGTACTAAGAGGTTTGATAACCCTGTCCTACCATTTAATG | target |
| AM161_SARS_COV2_S_D80_BETA | GGGTCTCTGGGACCAATGGTACTAAGAGGTTTGCTAACCCTGTCCTACCATTTAATG | target |
| AM164_SARS_COV2_S_L452_WT | GGATTCTAAGGTTGGTGGTAATTATAATTACCTGTATAGATTGTTTAGGAAGTCTAA | target |
| AM165_SARS_COV2_S_L452_DELTA | GGATTCTAAGGTTGGTGGTAATTATAATTACCGGTATAGATTGTTTAGGAAGTCTAA | target |
| AM168_SARS_COV2_S_S477_WT | GGTTTCAACTGAAATCTATCAGGCCGGTAGCACACCTTGTAATGGTGTTGAAGGTTT | target |
| AM169_SARS_COV2_S_S477_OMICRON | GGTTTCAACTGAAATCTATCAGGCCGGTAACAAACCTTGTAATGGTGTTGAAGGTTT | target |
| AM162_WT_crRNA_S_D80_nt | GGACCACCCCAAAAAUGAAGGGGACUAAAACGGGUUAUCAAACCUCUUAGU | crRNA |
| AM163_beta_crRNA_S_D80_20nt | GGACCACCCCAAAAAUGAAGGGGACUAAAACGGGUUAGCAAACCUCUUAGU | crRNA |
| AM166_WT_crRNA_L452_S_20nt | GGACCACCCCAAAAAUGAAGGGGACUAAAACCUAUACAGGUAAUUAUAAUU | crRNA |
| AM167_Delta_crRNA_L452R_S_20nt | GGACCACCCCAAAAAUGAAGGGGACUAAAACCUAUACCGGUAAUUAUAAUU | crRNA |
| AM170_WT_crRNA_S477_S_20nt | GGACCACCCCAAAAAUGAAGGGGACUAAAACCAAGGUGUGCUACCGGCCUG | crRNA |
| AM171_omicron_crRNA_477_478_S_20nt | GGACCACCCCAAAAAUGAAGGGGACUAAAACCAAGGUUUGUUACCGGCCUG | crRNA |

| | | |
|---|---|---|
| AM194_WT_crRNA_S_D80_20nt_MM19 | GGACCACCCCAAAAAUGAAGGGGACUAAAACGGGUUAUCAAACCUCUUACU | crRNA |
| AM195_beta_crRNA_S_D80_20nt_MM7_19 | GGACCACCCCAAAAAUGAAGGGGACUAAAACGGGUUAGCAAACCUCUUACU | crRNA |
| AM196_WT_crRNA_L452_S_20nt_MM19 | GGACCACCCCAAAAAUGAAGGGGACUAAAACCUAUACAGGUAAUUAUAAAU | crRNA |
| AM197_Delta_crRNA_L452R_S_20nt_MM7_19 | GGACCACCCCAAAAAUGAAGGGGACUAAAACCUAUACCGGUAAUUAUAAAU | crRNA |
| AM198_WT_crRNA_S477_S_20nt_MM19 | GGACCACCCCAAAAAUGAAGGGGACUAAAACCAAGGUGUGCUACCGGCCAG | crRNA |
| AM199_omicron_crRNA_477_478_S_20nt_MM7_19 | GGACCACCCCAAAAAUGAAGGGGACUAAAACCAAGGUUUGUUACCGGCCAG | crRNA |
| AM202_wt_crRNA_D80_v2 | GGACCACCCCAAAAAUGAAGGGGACUAAAACAAUGGUAGGACAGGGUUAUC | crRNA |
| AM203_beta_crRNA_D80_v2_MM19 | GGACCACCCCAAAAAUGAAGGGGACUAAAACAAUGGUAGGACAGGGUUAGC | crRNA |
| AM204_wt_crRNA_L452_v2 | GGACCACCCCAAAAAUGAAGGGGACUAAAACUUCCUAAACAAUCUAUACAG | crRNA |
| AM205_delta_crRNA_L452_v2_MM19 | GGACCACCCCAAAAAUGAAGGGGACUAAAACUUCCUAAACAAUCUAUACCG | crRNA |
| AM206_wt_crRNA_S477_v2 | GGACCACCCCAAAAAUGAAGGGGACUAAAACACACCAUUACAAGGUGUGCU | crRNA |
| AM207_omicron_crRNA_S477_v2_MM19 | GGACCACCCCAAAAAUGAAGGGGACUAAAACACACCAUUACAAGGUUUGUU | crRNA |
| AM212_WT_del_crRNA_S_D80_nt | GGCCACCCCAAAAAUGAAGGGGACUAAAACGGGUUAUCAAACCUCUUAGU | crRNA |
| AM213_beta_del_crRNA_S_D80_20nt | GGCCACCCCAAAAAUGAAGGGGACUAAAACGGGUUAGCAAACCUCUUAGU | crRNA |
| AM214_WT_del_crRNA_L452_S_20nt | GGCCACCCCAAAAAUGAAGGGGACUAAAACCUAUACAGGUAAUUAUAAUU | crRNA |
| AM215_Delta_del_crRNA_L452R_S_20nt | GGCCACCCCAAAAAUGAAGGGGACUAAAACCUAUACCGGUAAUUAUAAUU | crRNA |

| | | |
|---|---|---|
| AM216_WT_del_crRNA_S477_S_20nt | GGCCACCCCAAAAAUGAAGGGGACUAAAACCAAGGUGUGCUACCGGCCUG | crRNA |
| AM217_omicron_del_crRNA_477_478_S_20nt | GGCCACCCCAAAAAUGAAGGGGACUAAAACCAAGGUUUGUUACCGGCCUG | crRNA |
| AM222_crRNA_S_D80_20nt_MM19_mut_DR | GGCCACCCCAAAAAUGAAGGGGACUAAAACGGGUUAUCAAACCUCUUACU | crRNA |
| AM223_beta_crRNA_S_D80_20nt_MM7_19_mut_DR | GGCCACCCCAAAAAUGAAGGGGACUAAAACGGGUUAGCAAACCUCUUACU | crRNA |
| AM224_WT_crRNA_L452_S_20nt_MM19_mut_DR | GGCCACCCCAAAAAUGAAGGGGACUAAAACCUAUACAGGUAAUUAUAAAU | crRNA |
| AM225_Delta_crRNA_L452R_S_20nt_MM7_19_mut_DR | GGCCACCCCAAAAAUGAAGGGGACUAAAACCUAUACCGGUAAUUAUAAAU | crRNA |
| AM226_WT_crRNA_S477_S_20nt_MM19_mut_DR | GGCCACCCCAAAAAUGAAGGGGACUAAAACCAAGGUGUGCUACCGGCCAG | crRNA |
| AM227_omicron_crRNA_477_478_S_20nt_MM7_19_mut_DR | GGCCACCCCAAAAAUGAAGGGGACUAAAACCAAGGUUUGUUACCGGCCAG | crRNA |
| AM228_Liu_mut_DR_Lbu_20_pos7_G | GGCCACCCCAAAAAUGAAGGGGACUAAAACACAAAUGUAUCUGAAUAAAC | crRNA |
| AM229_Liu_mut_DR_Lbu_20_pos7_A | GGCCACCCCAAAAAUGAAGGGGACUAAAACACAAAUAUAUCUGAAUAAAC | crRNA |
| AM230_Liu_mut_DR_Lbu_20_pos7_U | GGCCACCCCAAAAAUGAAGGGGACUAAAACACAAAUUUAUCUGAAUAAAC | crRNA |
| AM243_WT_del_crRNA_S_D80_20nt_MM6 | GGCCACCCCAAAAAUGAAGGGGACUAAAACGGGUUUUCAAACCUCUUAGU | crRNA |
| AM244_beta_del_crRNA_S_D80_20nt_MM6 | GGCCACCCCAAAAAUGAAGGGGACUAAAACGGGUUUGCAAACCUCUUAGU | crRNA |
| AM245_WT_del_crRNA_L452_S_20nt_MM6 | GGCCACCCCAAAAAUGAAGGGGACUAAAACCUAUAGAGGUAAUUAUAAUU | crRNA |
| AM246_Delta_del_crRNA_L452R_S_20nt_MM6 | GGCCACCCCAAAAAUGAAGGGGACUAAAACCUAUAGCGGUAAUUAUAAUU | crRNA |
| AM247_WT_del_crRNA_S477_S_20nt_MM6 | GGCCACCCCAAAAAUGAAGGGGACUAAAACCAAGGAGUGCUACCGGCCUG | crRNA |

| | | |
|---|---|---|
| AM248_omicron_del_crRNA_477_478_S_20nt_MM6 | GGCCACCCCAAAAAUGAAGGGGACUAAAACCAAGGAUUGUUACCGGCCUG | crRNA |
| AM249_WT_crRNA_S_D80_20nt_MM6_19 | GGACCACCCCAAAAAUGAAGGGGACUAAAACGGGUUUUCAAACCUCUUACU | crRNA |
| Liu_modified_localGC_3_G | GGGAAACCAA GAAUAAGAGCUCACUCGGAUACAUUUGC CACAGCAGAAGCCCACAC | target |
| Liu_modified_localGC_3_C_MM | GGGAAACCAA GAAUAAGAGCUCACUCGGAUAgAUUUGC CACAGCAGAAGCCCACAC | target |
| Liu_modified_totalGC_50_G | GGGAAACCAA GAAGAAGAAUUUAUUUAGAUGCGCUUAU CACAGCAGAAGCCCACAC | target |
| Liu_modified_totalGC_50_C_MM | GGGAAACCAA GAAGAAGAAUUUAUUUAGAUGgGCUUAU CACAGCAGAAGCCCACAC | target |
| AM254_Lbu_mut_DR_modified_localGC_3_C | GGCCACCCCAAAAAUGAAGGGGACUAAAACAUAAGCCCAUCUAAAUAAAU | crRNA |
| AM255_Lbu_mut_DR_modified_localGC_3_G | GGCCACCCCAAAAAUGAAGGGGACUAAAACAUAAGCGCAUCUAAAUAAAU | crRNA |
| AM256_Lbu_mut_DR_modified_totalGC_50_C | GGCCACCCCAAAAAUGAAGGGGACUAAAACGCAAAUCUAUCCGAGUGAGC | crRNA |
| AM257_Lbu_mut_DR_modified_totalGC_50_G | GGCCACCCCAAAAAUGAAGGGGACUAAAACGCAAAUGUAUCCGAGUGAGC | crRNA |
| AM261_WT_del_crRNA_L452R_S_20nt_MM19 | GGCCACCCCAAAAAUGAAGGGGACUAAAACUUCCUAAACAAUCUAUACAG | crRNA |
| AM262_Delta_del_crRNA_L452R_S_20nt_MM19 | GGCCACCCCAAAAAUGAAGGGGACUAAAACUUCCUAAACAAUCUAUACCG | crRNA |

**Table S4. Oligonucleotides used for SARS-CoV-2 amplification.**

| Name | Sequence 5′-3′ | Ref. |
|---|---|---|
| AM238_SARS_CoV2_D80A_fwd_Arizti-Sanz | *GAAATTAATACGACTCACTATAGGG* CAACTCAGGACTTGTTCTTACCTTTCTTTTCC | (13) |
| AM239_SARS_CoV2_D80A _rev_Arizti-Sanz | AAGCAAAATAAACACCATCATTAAAT | (13) |
| AM263_RPA_fwd_Yang_SARS_S452_T7 | *GAAATTAATACGACTCACTATAGGG* CTTGATTCTAAGGTTGGTGGTAATTATAAT | (14) |
| AM264_RPA_rev_Yang_SARS_S452 | AAGGTTTGAGATTAGACTTCCTAAACAATC | (14) |
| AM220_SARS_CoV2_T478K_fwd_Yang | *GAAATTAATACGACTCACTATAGGG* TTGAGAGAGATATTTCAACTGAAATCTATC | (14) |
| AM221_ SARS_CoV2_T478K_rev_Yang | AGTGGGTTGGAAACCATATGATTGTAAAGG | (14) |

**Table S5. Nomenclature of the SARS-CoV-2 lineages referenced, and viral genomes used in this study.**

| This study | WHO classification | PANGO lineage | GISAID accessions of virus used in this study | Viral isolate, BEI repository number |
|---|---|---|---|---|
| Ancestral | - | A | EPI_ISL_412028 | Hong Kong/VM20001061/2020, NR-52282 |
| Beta | Beta | B.1.351 | EPI_ISL_678570<br>EPI_ISL_678615 | South Africa/KRISP-K005325/2020, NR-54009 |
| Delta | Delta | B.1617.2 | N/A* | USA/PHC658/2021, NR-55611 |
| Omicron | Omicron | B.1.1.529 | EPI_ISL_7160424 | USA/MD-HP20874/2021, NR-56461 |

* Sequence information can be found in the BEI repository for hCoV-19/USA/PHC658/2021 (NR-55611)

**Table S6. RT-qPCR values of SARS-CoV-2 strains used in this study.**

| Viral strain | 2019-nCoV N1 Ct mean value | Ct standard deviation | 2019-nCoV N2 Ct mean value | Ct standard deviation |
|---|---|---|---|---|
| Hong Kong/VM20001061/2020 | 16.975 | 0.050 | 15.211 | 0.346 |
| South Africa/KRISP-K005325/2020 | 14.209 | 0.080 | 13.464 | 0.569 |
| USA/PHC658/2021 | 21.040 | 0.076 | 19.952 | 0.384 |
| USA/MD-HP20874/2021 | 17.939 | 0.071 | 15.780 | 0.398 |

**Supplementary Text 1. Confirmation of correct mismatched base conformation in our simulation**

To verify the conformation of the RNA bases at the mismatched positions in our simulations, we performed enhanced sampling simulations, using the Umbrella Sampling (US) method (8). We systematically explored the dihedral angle $\theta$, defining the flipping of the mismatched base relative to the preceding matched base pair (**Figure S6**, see *Supplementary Materials and Method*). This enabled computing the free energy profiles corresponding to the dihedral scans and identifying the conformations associated with the energetic minima. The free energy profiles of the systems including a single mismatch at the spacer nucleotide positions 4, 7, or 11 (**Figure S6B**) were compared with the distributions of the $\theta$ dihedrals sampled by classical MD simulations (**Figure S6C**). We observe an excellent overlap between the free energy minima and the peaks of the distributions computed from classical MD simulations. This agreement strengthens the calculations of the SNR profiles through classical MD simulations since the conformations of single mismatches correspond to the preferred base conformation. The convergence of the US simulations is also reported in **Figure S6D**.

**Supplementary Text 2. LbuCas13a discrimination assays of SARS-CoV-2-like short RNAs.**

To test different strategies suitable for SARS-CoV-2 discrimination, we synthesized 57-nucleotide long RNAs that contain a genomic viral sequence of interest (S gene), and their SNPs depending on a different variant (Beta, Delta, Omicron). We tested Cas13a-activation with the different LbuCas13a variants and crRNAs described previously against these short RNAs at 10 nM final concentration.

Given that mismatches at positions 7 and 19 – relative to the crRNA yielded significant decrease in nuclease activation, we designed crRNA combinations where the SNP would occur at these positions. Using a mismatch at position 7 only resulted in limited discrimination for LbuCas13a$^{R377A}$ and was not generalizable for every sample combination tested (**Figure S9A**), while other variants did not yield discrimination with this design (**Figure S9A**). The presence of the SNP-detecting mismatch at position 19 resulted in about 50% decrease in activity for

LbuCas13a$^{R377A}$ and for most crRNA: target RNA combinations tested it resulted in clear discrimination (**Figure S9B**), whereas effects on LbuCas13a$^{N378A}$ discrimination ability were modest (**Figure S9B**).

Consequently, we tested an approach in which we primed the enzyme with an initial synthetic mismatch at position 19 for all cases and then used position 7 for SNP detection. The decision to use those two positions is based on the high sensitivity we observed. Similar strategies have been described for Cas13-based detection platforms (15,16). Our approach resulted in strong loss of activity for LbuCas13a$^{R377A}$ (**Figure S9C**) that might render this combination less desirable for SNP-detection goals. Importantly, introducing this synthetic mismatch yielded substantial discrimination in LbuCas13a$^{N378A}$, as shown by strong reduction in activity for most of the mismatched samples at the concentration tested of 10 nM target RNA (**Figure S9C**). This is particularly exciting as a combined use of variant LbuCas13a$^{N378A}$ primed with an initial synthetic mismatch could readily be deployed for SNP detection. These results suggest this dual mismatch approach is very promising for the development of novel high-fidelity Cas13 SNP diagnostics. Nevertheless, for the SARS-CoV-2 D80A assay design, the discrimination achieved with the ancestral crRNA - VOC target was suboptimal, which could increase the likelihood of a false positive, underscoring mismatch base-pair orientation differences and the need to carefully test Cas13 crRNA design if SNP-discrimination is desired.

Additionally, we attempted to use the same crRNA design with the discriminatory position at the 7th nucleotide, but with the truncated direct repeat and viral variant discrimination could not be achieved. For LbuCas13a$^{R377A}$ activity was impaired even for perfect matched target RNAs, suggesting the combination is detrimental for overall activity. For the other variants and wild-type Cas13, no significant difference in apparent nuclease activation could be appreciated for mismatched crRNA-target pairs (**Figure S10A**).

The extreme sensitivity of mismatches at positions 7 and 19 when using a truncated direct repeat (**Figure 4D-E**) could not be productively combined and harnessed for SNP discrimination of these SARS-CoV-2 mutants as all Cas13 variants displayed compared to the wild-type Cas13

(**Figure S10B**), as in most cases cleavage activity was too weak to be desirable for diagnostic applications.

It is possible that combining two positions in the crRNA-target duplex that are particularly sensitive can easily overcome the balance between robust activation and sensitive SNP-detection (low sensitivity), thus rendering this strategy less desirable for widespread application and diagnostic purposes. Therefore, we finally tested SNP-discrimination power by keeping the most sensitive position (the 7th nucleotide) and introducing a synthetic mismatch adjacent to it (6th nucleotide) which by itself does not alter the assay's sensitivity in our previous assays (**Figure 4E-F**). We performed assays against the three SARS-CoV-2 regions of interest in the S gene using our truncated crRNAs and we observed strong activity with WT LbuCas13a, no substantial activity with LbuCas13a$^{R377A}$ (except in two instances) (**Figure S10C**). With LbuCas13a$^{N378A}$ and, less extent, LbuCas13a$^{R973A}$, they showed almost no activation for some at least one mismatched crRNA-target pairs (~10% relative to WT Cas13 with perfect matched RNAs). Interestingly, the opposite mismatched pair for a given assay (either VOC crRNA vs. ancestral target or ancestral crRNA vs. VOC target, depending on the assay) did not result in markedly less nuclease activity compared to perfect matched crRNA-target pairs for that particular design (**Figure S10C**). Taken together, we showed that combining specific Cas13 variants with direct-repeat truncation in the crRNA with a synthetic mismatch at position 6 and employing position 7 of the crRNA spacer for SNP discrimination can be used.

In sum, the apparent resistance of these sequences to Cas13-based SNP discrimination might be due to sequence-specific effects, for example, the type of mismatch and the orientation of the base pairs that participate in the mismatch, as the nucleotide changes that arise naturally in these viral variants are different to the mismatches we introduced (for example, for the SARS-CoV-2 spike:D80A variant, there is an A-to-C mutation). This would suggest that additional strategies might need to be followed to overcome the diversity in sequences that might be encountered when designing new assays for SNP-detection.

### Supplementary Figures

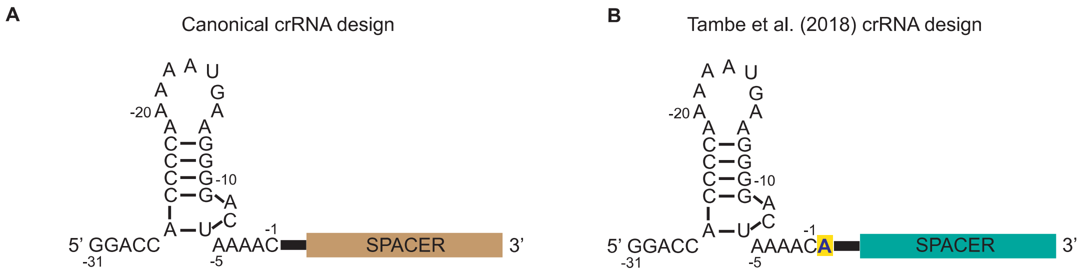


**Figure S1 | Differences in previously published single-mismatch profiling can be accounted to alternative crRNA designs.** Schematics comparing a canonical crRNA design **A.** as used in this study and **B.** the one used by Tambe et al. (2018) where the direct repeat ends with an additional adenine.

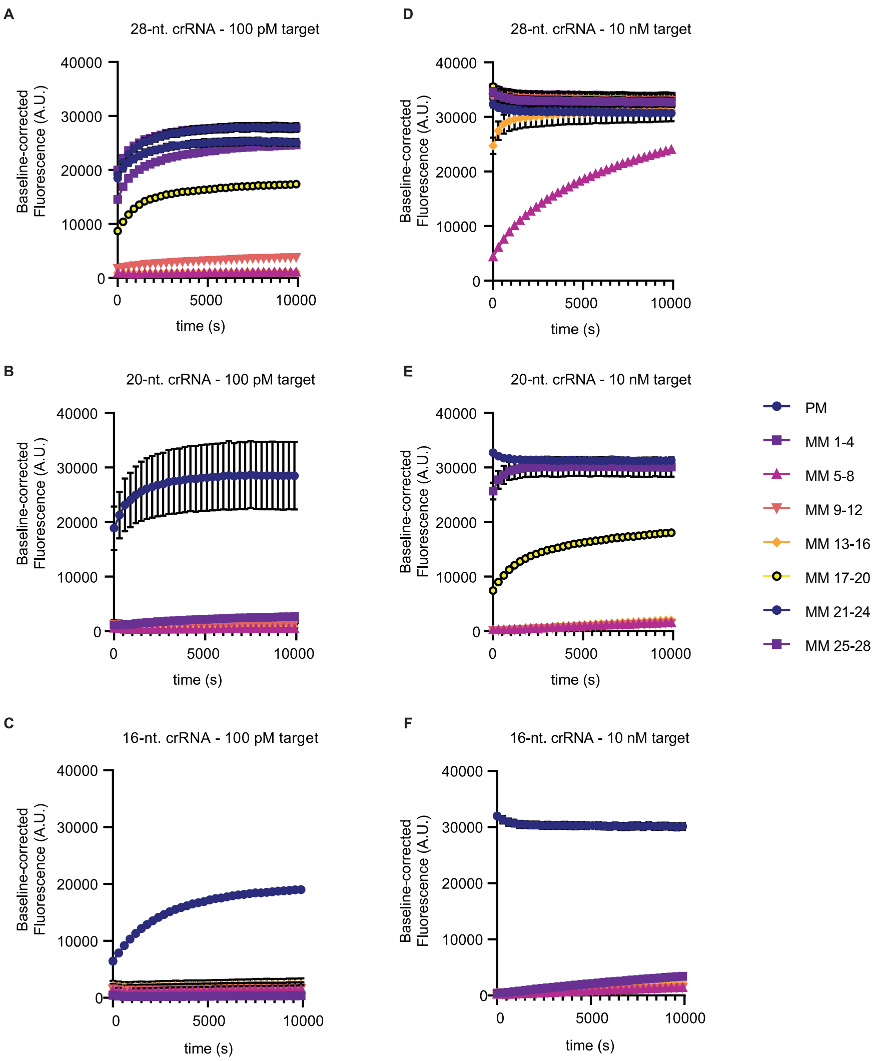


**Figure S2 | Related to Figure 1. LbuCas13a reporter cleavage time-course for each 20 and 28 nucleotide spacer lengths against the same target that are a perfect match (PM) or contain four consecutive mismatches (MM) across the crRNA:spacer.** These correspond to the experiments shown in Fig 1 E and F, as follows: **A.** 28 nucleotide spacer and with 100 pM RNA target final concentration; **B.** 20 nucleotide spacer and with 100 pM RNA target final concentration; **C.** 16 nucleotide spacer and with 100 pM RNA target final concentration **D.** 28 nucleotide spacer and with 10 nM RNA target final concentration; **E.** 20 nucleotide spacer and with 10 nM RNA target final concentration; **F.** 16 nucleotide spacer and with 10 nM RNA target final concentration. PM: perfect match RNA; MM: mismatches.

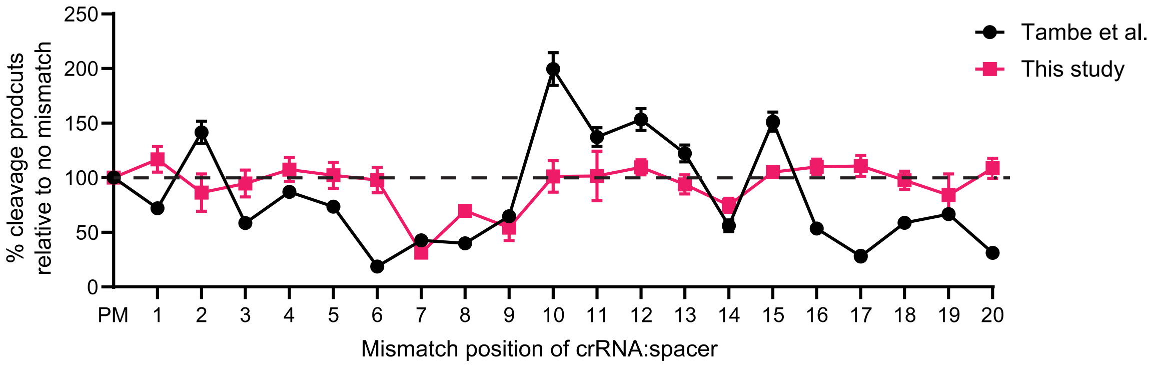


**Figure S3 | Related to Figure 2.** Comparison of LbuCas13a activation against single nucleotide mismatches between this study and *Tambe et. al*. using 20 nucleotide spacers and 100 pM RNA target. PM: perfect match RNA.

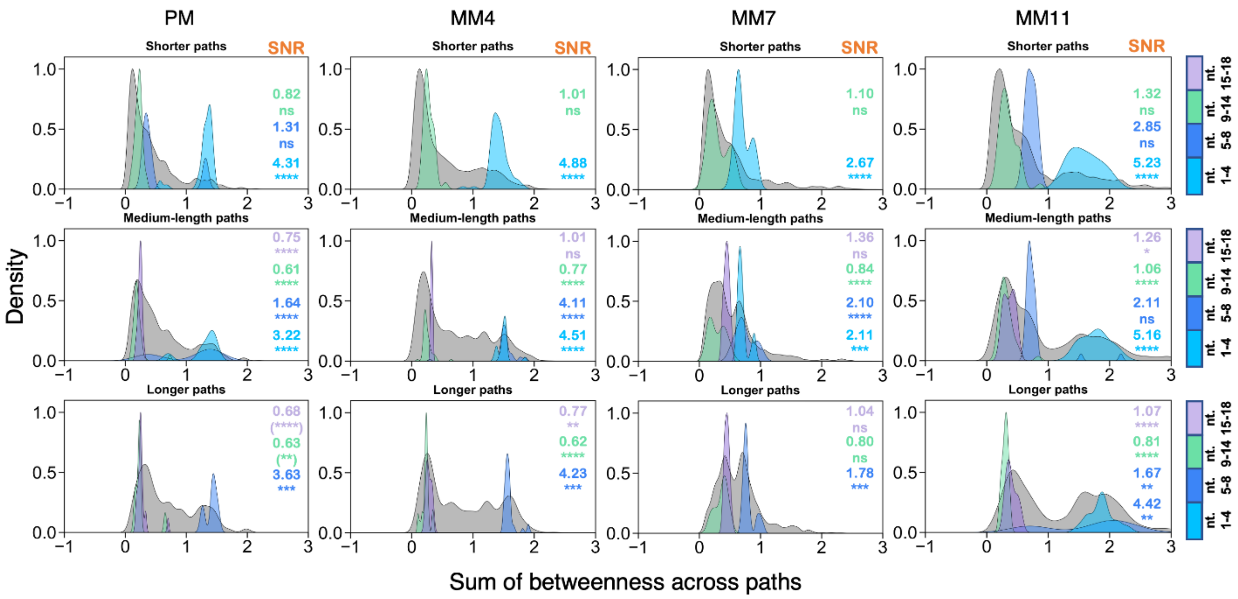


**Figure S4. Distribution of the signals over the noise in the tgRNA-bound Cas13 systems PM, MM4, MM7, and MM11.** Distribution of the signals from the crRNA spacer regions to the catalytic core residues, plotted on the background of noise (grey), across short (6-8 edge counts), medium (9-11), and long paths (12-14). The statistical significance of the signals over the noise was computed using z-score statistics with a two-tailed hypothesis (P-value reference: not significant, ns P > 0.01, * P ≤ 0.01, ** P ≤ 0.001, *** P ≤ 0.0001, **** P ≤ 0.00001). Values of the Signal-to-Noise Ratio (SNR) are also reported for each sourcing region.

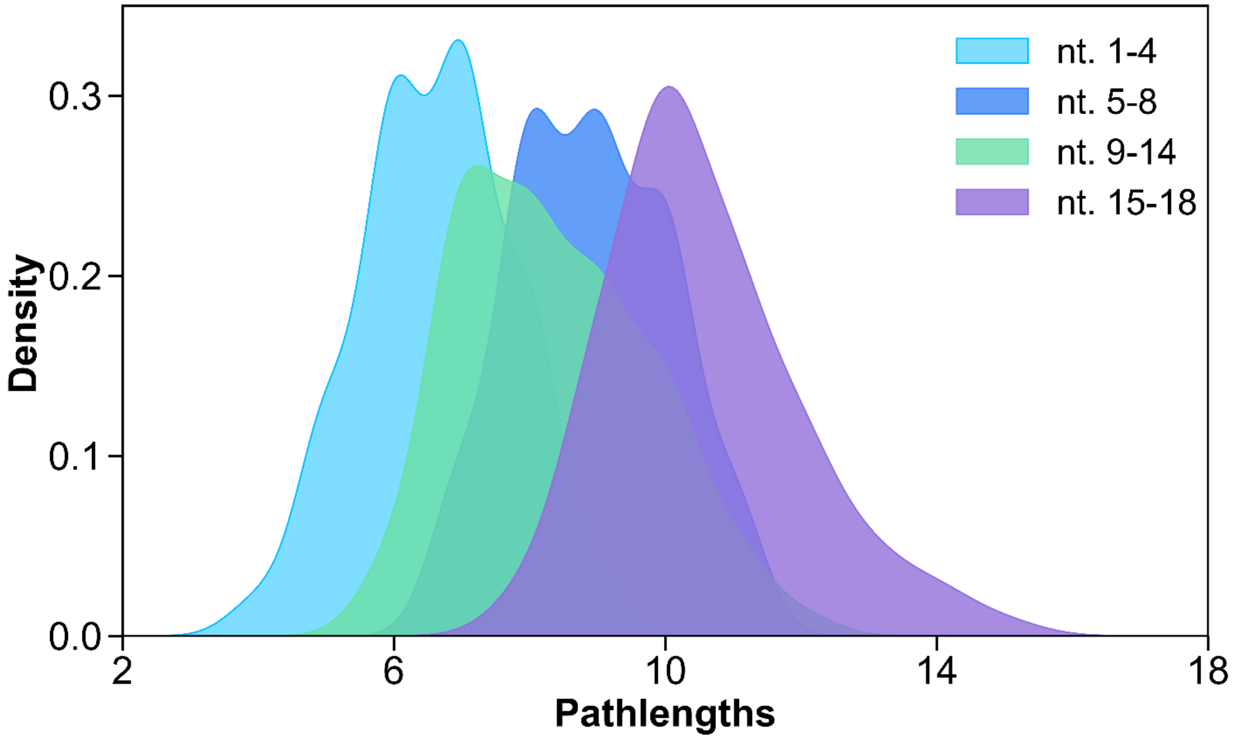


**Figure S5.** Kernel density estimation of pathlengths in terms of the number of edges connecting the crRNA spacer regions (i.e., nt. 1-4 (cyan), nt. 5-8 (blue), nt. 9-14 (green), and nt. 15-18 (violet)) to the catalytic core residues (R472, H1053, H477, R1048) in the perfectly matched Cas13a: crRNA: target-RNA complex. Pathlengths are calculated for the optimal and top five sub-optimal pathways.

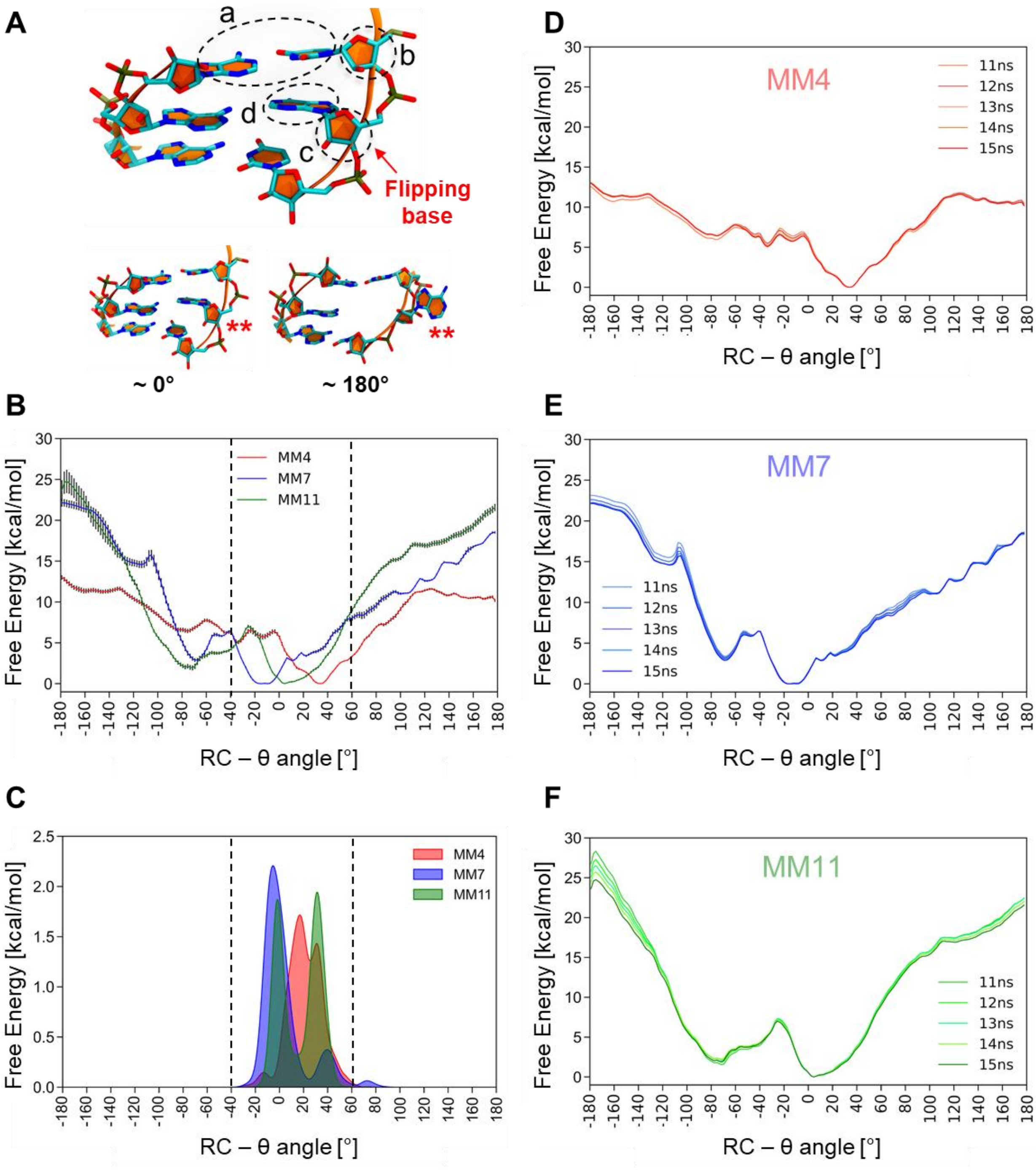


**Figure S6.** Free energy profiles describing base flipping at mismatched positions. **A**. Overview of the reaction coordinate (RC) used for free energy simulations. Upper panel: The pseudo dihedral angle θ describes the flipping of the mismatched base relative to the adjacent matched base pair, as previously used in computational studies of base flipping (9,10). The θ dihedral is computed between the centers of mass (COM) of four groups of atoms: (a) the heavy atoms of the 5′ base-pair adjacent to the flipping base, (b) the sugar moiety of the adjacent 5′ base, (c) the sugar moiety of the flipping base, and (d) the flipping base. Lower panel: Conformation of the flipping base (**) at θ ~0° and ~180°. **B**. Free energy profiles for base flipping at position 4 (MM4), 7 (MM7), or 11 (MM11), computed along θ using Umbrella Sampling (US) simulations. Vertical dashed lines indicate the dihedral range corresponding to the conformations associated with the energetic minima. Error bars represent the standard deviation obtained from the free energy profiles, computed from the last 5 ns and at 1 ns intervals at each point of the RC. **C**. Distribution of the θ dihedral angle sampled through classical MD simulations of the single mismatched Cas13a systems. **D-F**. Convergence of the free energy profiles, computed from the last 5 ns and at 1 ns intervals for each US window.

**A**

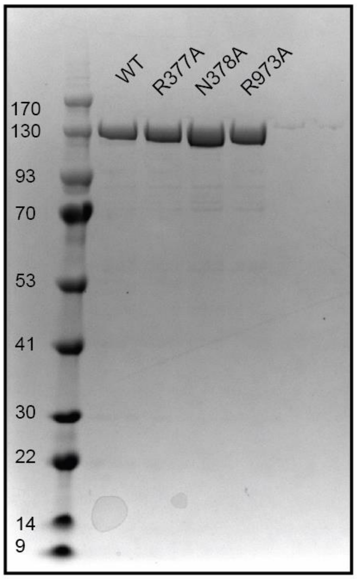


**B**

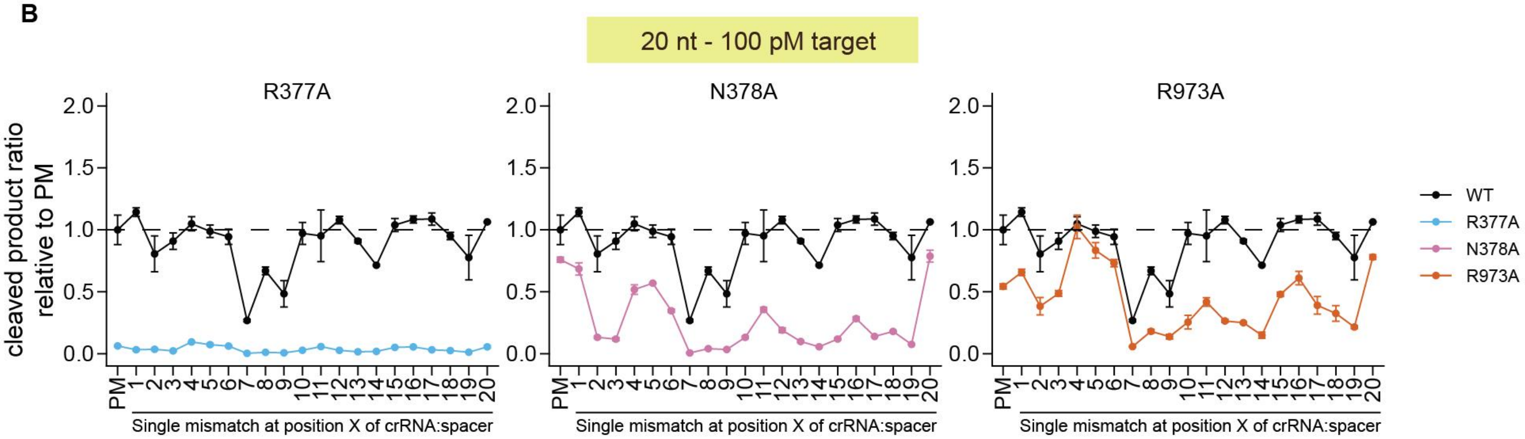


**Figure S7 | Related to Figure 3. A.** SDS-PAGE gel showing ~2 ug of purified wild-type (WT) LbuCas13a and variants R377A, N378A, and R973A by Coomassie staining. **B.** Single-nucleotide mismatch profiling of LbuCas13a variants with 20 nucleotide spacers and 100 pM RNA target. Results are normalized and represented by ratio of cleavage product compared to wild-type (WT) LbuCas13a targeting a non-mismatched RNA (PM: perfect match).

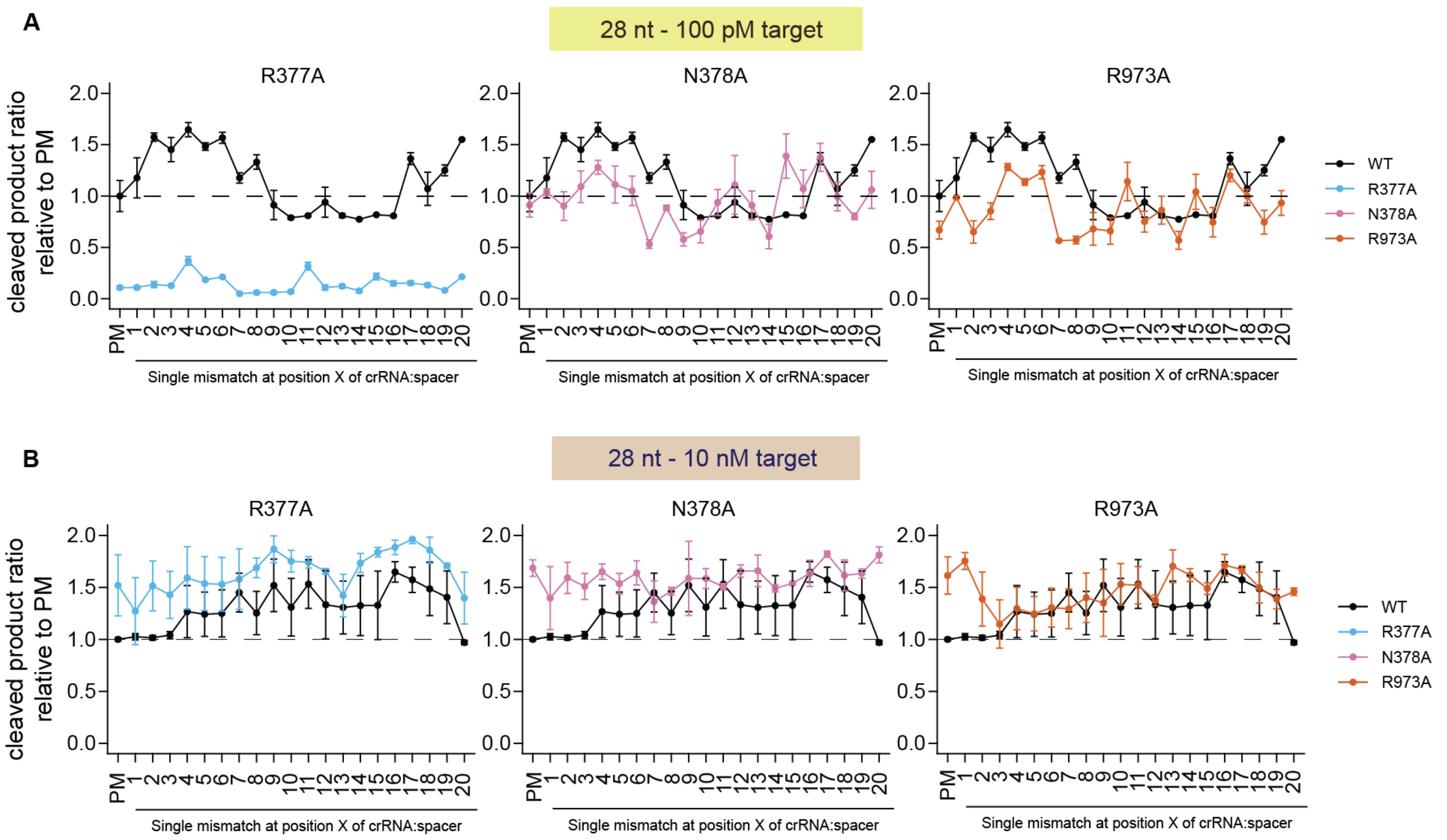


**Figure S8 | Related to Figure 3.** Single-nucleotide mismatch profiling of LbuCas13a variants with 28 nucleotide spacers and **A.** 100 pM RNA target or **B.** 10 nM RNA target. Results are normalized and represented by ratio of cleavage product compared to wild-type (WT) LbuCas13a targeting a non-mismatched RNA (PM: perfect match).

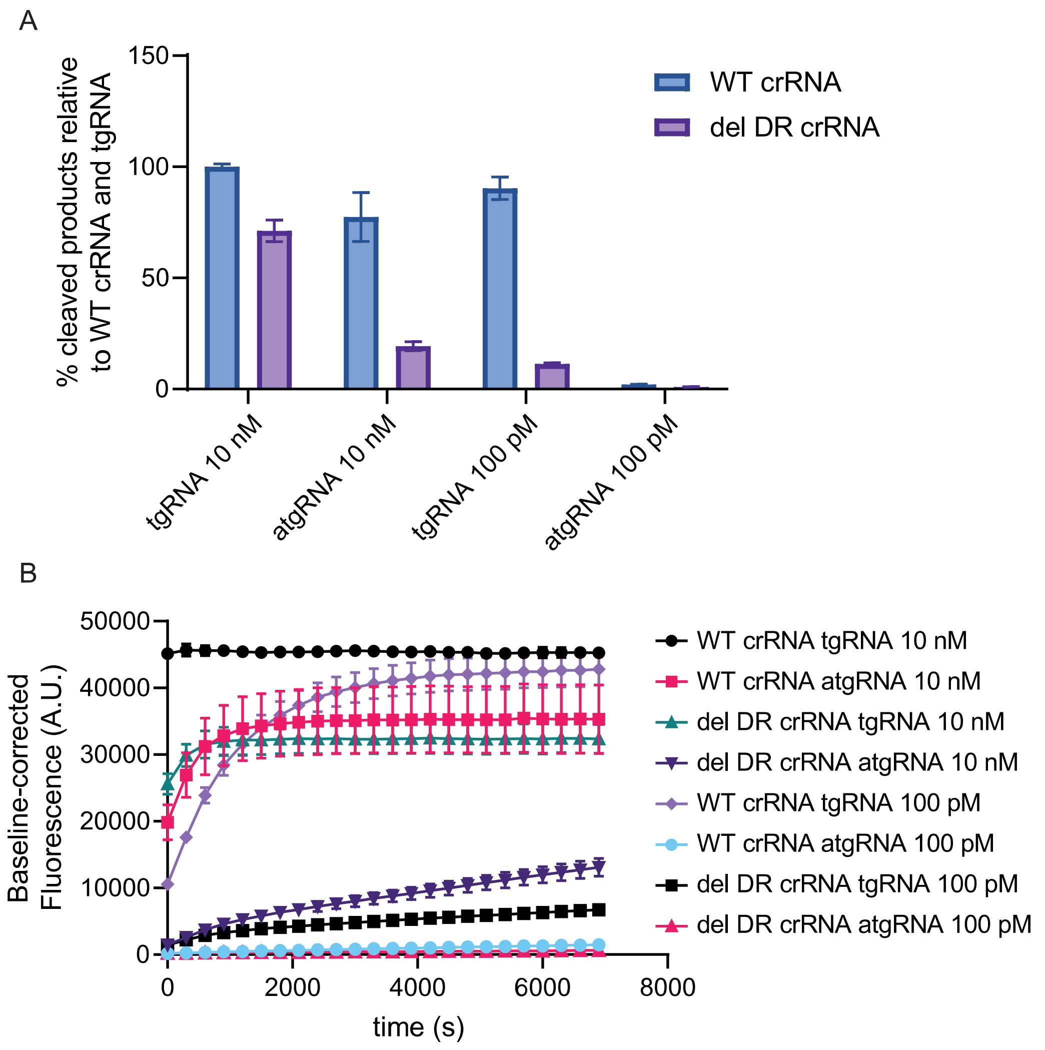


**Figure S9 | Related to Figure 4. Restoring crRNA direct repeat to full length results in increased LbuCas13a cleavage rates and higher tolerance to atgRNAs**. A. End-point background corrected fluorescence values at one-hour from RNA cleavage experiments by WT LbuCas13a by the addition of 100 pM or 10 nM tgRNA or atgRNA for spacer used by *Meeske et al.* (2018). Normalized as percent cleavage product generated relative to WT LbuCas13a with tgRNA. **B.** Two-hour time course of background corrected fluorescence measurements from RNA cleavage experiments by WT LbuCas13a by the addition of 100 pM or 10 nM tgRNA or atgRNA for spacer used by *Meeske et al.* (2018). *WT crRNA*: full length crRNA; *del DR crRNA*: crRNA with deletion of $A_{-29}$ in the direct repeat of the crRNA.

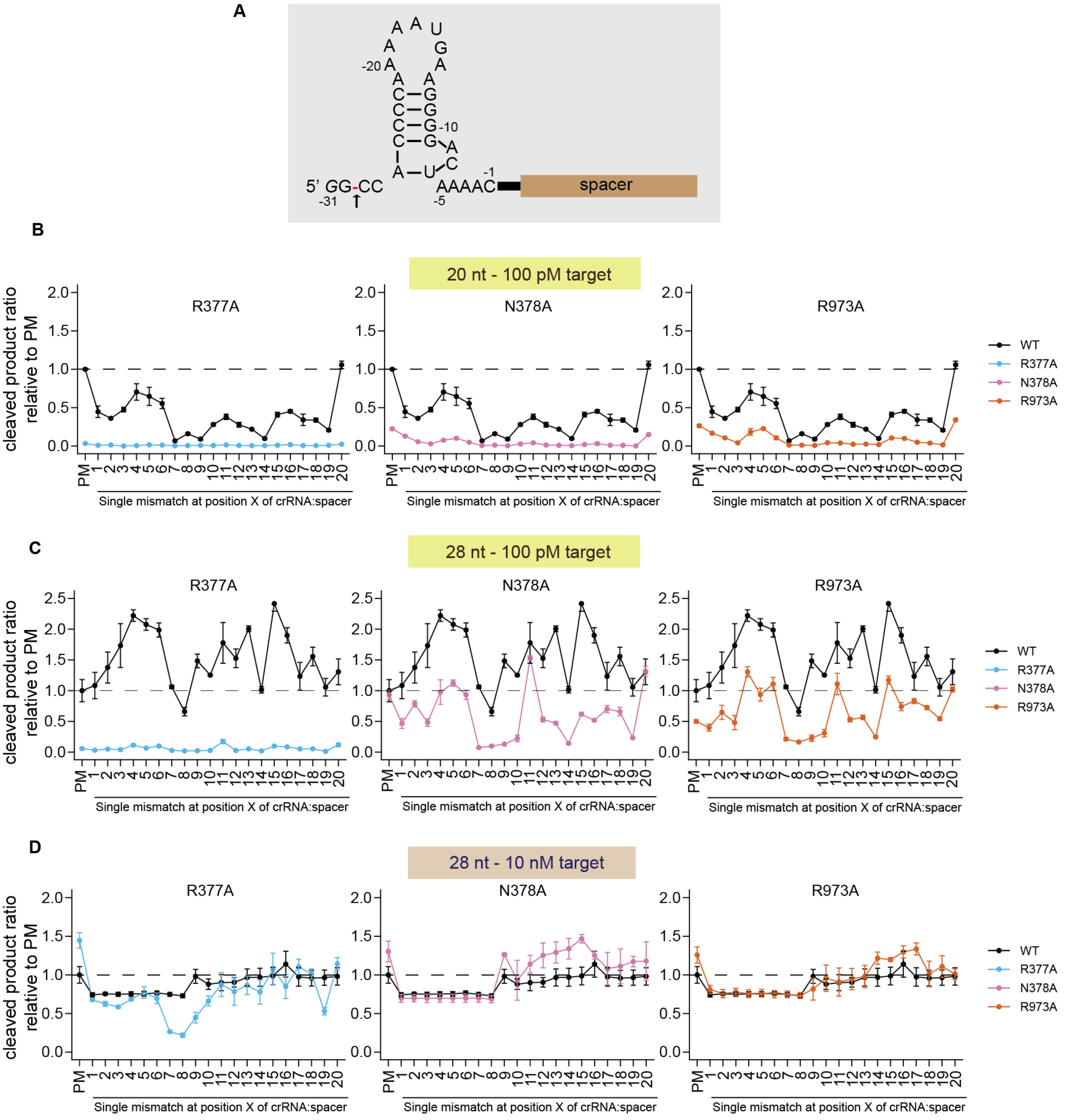
A
5' GG-CC
-31
-20
-10
-5
-1
AAAAC
spacer
B
20 nt - 100 pM target
R377A
N378A
R973A
cleaved product ratio relative to PM
Single mismatch at position X of crRNA:spacer
WT
R377A
N378A
R973A
C
28 nt - 100 pM target
D
28 nt - 10 nM target

**Figure S10 | Related to Figure 4. A.** crRNA design of the truncated direct repeat, as reported in *Meeske et. al* (2018). **B-D.** Single-nucleotide mismatch profiling of LbuCas13a variants with **B.** 20 nucleotide spacers and 100 pM RNA activator, **C.** 28 nucleotide spacers with 100 pM target or **D.** 28 nucleotide spacers with 10 nM target. Results are normalized and represented by ratio of cleavage product compared to wild-type (WT) LbuCas13a targeting a non-mismatched RNA (PM: perfect match).

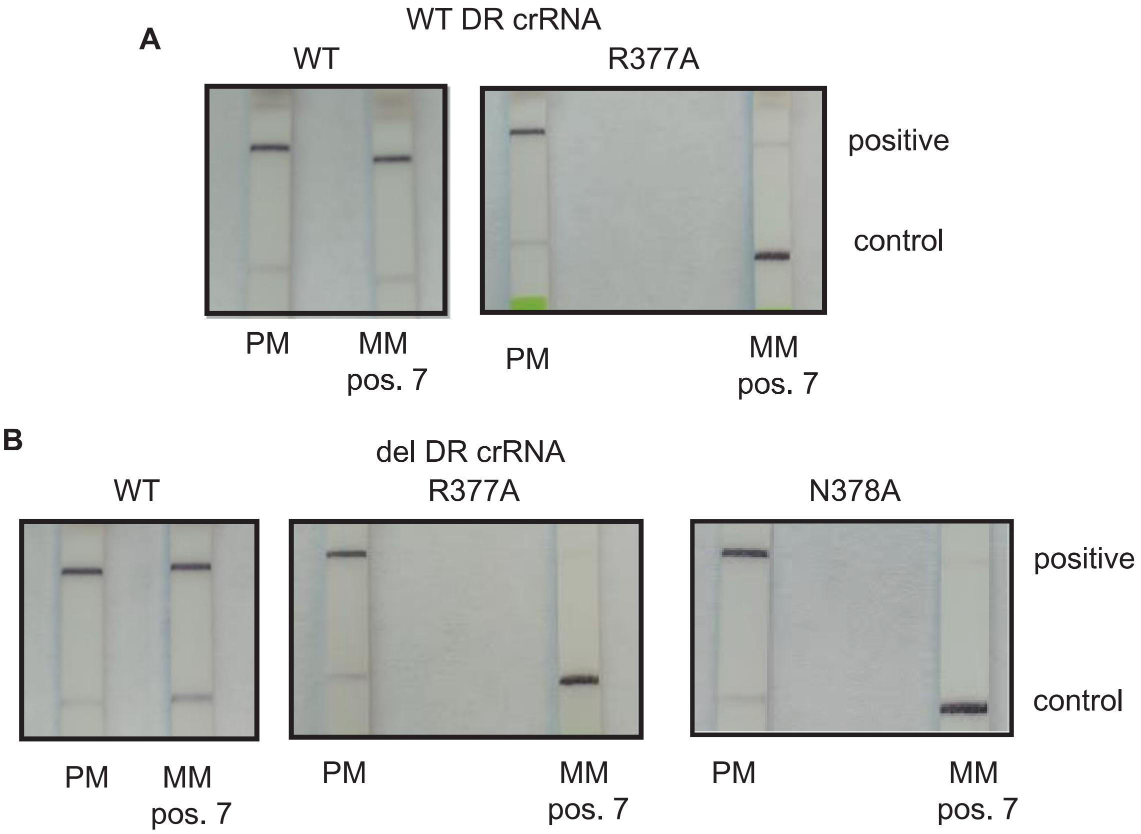


**Figure 11 | Related to Figures 3 and 4.** Lateral flow readout after 1 hour of LbuCas13a WT or selected variants reporter cleavage in the presence of short RNA targets for single-nucleotide discrimination. **A.** Comparison between WT LbuCas13a and LbuCas13a$^{R377A}$ with a full direct repeat crRNA lateral flow readout detecting perfect-matched RNA target (PM) or with a mismatch at position 7 (MM pos. 7), relative to 5′ of the spacer. **B.** Comparison between WT LbuCas13a, LbuCas13a$^{R377A}$ and LbuCas13a$^{N378A}$ with a truncated direct repeat crRNA lateral flow readout detecting perfect-matched RNA target (PM) or with a mismatch at position 7 (MM pos. 7), relative to 5′ of the spacer.

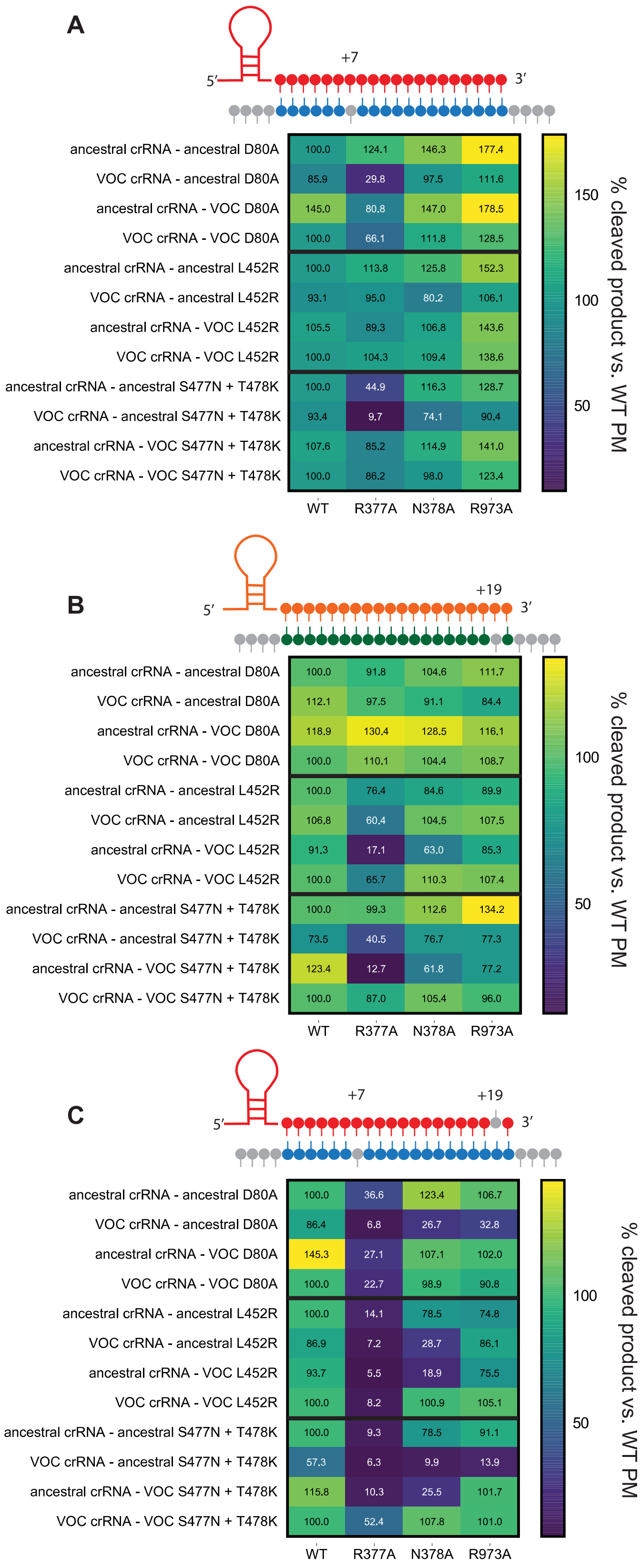
A
+7
5′
3′
ancestral crRNA - ancestral D80A 100.0 124.1 146.3 177.4
VOC crRNA - ancestral D80A 85.9 29.8 97.5 111.6
ancestral crRNA - VOC D80A 145.0 80.8 147.0 178.5
VOC crRNA - VOC D80A 100.0 66.1 111.8 128.5
ancestral crRNA - ancestral L452R 100.0 113.8 125.8 152.3
VOC crRNA - ancestral L452R 93.1 95.0 80.2 106.1
ancestral crRNA - VOC L452R 105.5 89.3 106.8 143.6
VOC crRNA - VOC L452R 100.0 104.3 109.4 138.6
ancestral crRNA - ancestral S477N + T478K 100.0 44.9 116.3 128.7
VOC crRNA - ancestral S477N + T478K 93.4 9.7 74.1 90.4
ancestral crRNA - VOC S477N + T478K 107.6 85.2 114.9 141.0
VOC crRNA - VOC S477N + T478K 100.0 86.2 98.0 123.4
WT R377A N378A R973A
150
100
50
% cleaved product vs. WT PM
B
+19
5′
3′
ancestral crRNA - ancestral D80A 100.0 91.8 104.6 111.7
VOC crRNA - ancestral D80A 112.1 97.5 91.1 84.4
ancestral crRNA - VOC D80A 118.9 130.4 128.5 116.1
VOC crRNA - VOC D80A 100.0 110.1 104.4 108.7
ancestral crRNA - ancestral L452R 100.0 76.4 84.6 89.9
VOC crRNA - ancestral L452R 106.8 60.4 104.5 107.5
ancestral crRNA - VOC L452R 91.3 17.1 63.0 85.3
VOC crRNA - VOC L452R 100.0 65.7 110.3 107.4
ancestral crRNA - ancestral S477N + T478K 100.0 99.3 112.6 134.2
VOC crRNA - ancestral S477N + T478K 73.5 40.5 76.7 77.3
ancestral crRNA - VOC S477N + T478K 123.4 12.7 61.8 77.2
VOC crRNA - VOC S477N + T478K 100.0 87.0 105.4 96.0
WT R377A N378A R973A
100
50
% cleaved product vs. WT PM
C
+7
+19
5′
3′
ancestral crRNA - ancestral D80A 100.0 36.6 123.4 106.7
VOC crRNA - ancestral D80A 86.4 6.8 26.7 32.8
ancestral crRNA - VOC D80A 145.3 27.1 107.1 102.0
VOC crRNA - VOC D80A 100.0 22.7 98.9 90.8
ancestral crRNA - ancestral L452R 100.0 14.1 78.5 74.8
VOC crRNA - ancestral L452R 86.9 7.2 28.7 86.1
ancestral crRNA - VOC L452R 93.7 5.5 18.9 75.5
VOC crRNA - VOC L452R 100.0 8.2 100.9 105.1
ancestral crRNA - ancestral S477N + T478K 100.0 9.3 78.5 91.1
VOC crRNA - ancestral S477N + T478K 57.3 6.3 9.9 13.9
ancestral crRNA - VOC S477N + T478K 115.8 10.3 25.5 101.7
VOC crRNA - VOC S477N + T478K 100.0 52.4 107.8 101.0
WT R377A N378A R973A
100
50
% cleaved product vs. WT PM

**Figure S12 | Related to Figure 5. A-C.** Heatmaps of percentage of cleaved product for each LbuCas13a variant when activated with a SARS-CoV-2 short RNA fragment, compared to wild-type (WT) LbuCas13a with a perfect match RNA (PM). Values assessed after 1 hour incubation using 10 nM of target RNA. crRN–A and target were either designed for the ancestral/Wuhan strain or to a given variant-of-concern (VOC) strain for the S gene as follows: beta (D80A), delta (L452R), omicron (S477N+T478K region). The different designs are **A.** SNP occurs at position 7 relative to crRNA; **B.** SNP occurs at position 19 relative to crRNA; **C.** enzyme is primed with a mismatch at position 19 in all cases and SNP occurs at position 7 relative to crRNA.

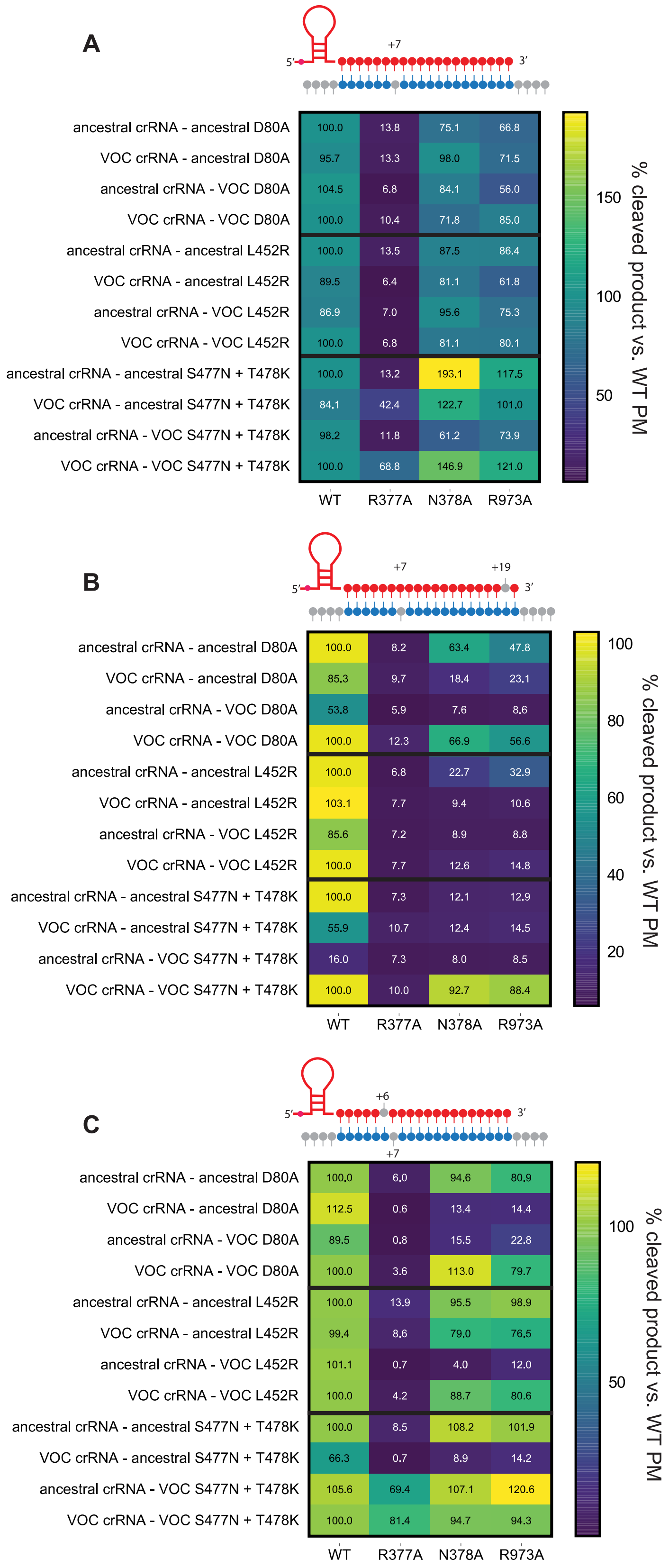
A
5′
+7
3′
ancestral crRNA - ancestral D80A 100.0 13.8 75.1 66.8
VOC crRNA - ancestral D80A 95.7 13.3 98.0 71.5
ancestral crRNA - VOC D80A 104.5 6.8 84.1 56.0
VOC crRNA - VOC D80A 100.0 10.4 71.8 85.0
ancestral crRNA - ancestral L452R 100.0 13.5 87.5 86.4
VOC crRNA - ancestral L452R 89.5 6.4 81.1 61.8
ancestral crRNA - VOC L452R 86.9 7.0 95.6 75.3
VOC crRNA - VOC L452R 100.0 6.8 81.1 80.1
ancestral crRNA - ancestral S477N + T478K 100.0 13.2 193.1 117.5
VOC crRNA - ancestral S477N + T478K 84.1 42.4 122.7 101.0
ancestral crRNA - VOC S477N + T478K 98.2 11.8 61.2 73.9
VOC crRNA - VOC S477N + T478K 100.0 68.8 146.9 121.0
WT R377A N378A R973A
150
100
50
% cleaved product vs. WT PM
B
5′
+7
+19
3′
ancestral crRNA - ancestral D80A 100.0 8.2 63.4 47.8
VOC crRNA - ancestral D80A 85.3 9.7 18.4 23.1
ancestral crRNA - VOC D80A 53.8 5.9 7.6 8.6
VOC crRNA - VOC D80A 100.0 12.3 66.9 56.6
ancestral crRNA - ancestral L452R 100.0 6.8 22.7 32.9
VOC crRNA - ancestral L452R 103.1 7.7 9.4 10.6
ancestral crRNA - VOC L452R 85.6 7.2 8.9 8.8
VOC crRNA - VOC L452R 100.0 7.7 12.6 14.8
ancestral crRNA - ancestral S477N + T478K 100.0 7.3 12.1 12.9
VOC crRNA - ancestral S477N + T478K 55.9 10.7 12.4 14.5
ancestral crRNA - VOC S477N + T478K 16.0 7.3 8.0 8.5
VOC crRNA - VOC S477N + T478K 100.0 10.0 92.7 88.4
WT R377A N378A R973A
100
80
60
40
20
% cleaved product vs. WT PM
C
5′
+6
3′
+7
ancestral crRNA - ancestral D80A 100.0 6.0 94.6 80.9
VOC crRNA - ancestral D80A 112.5 0.6 13.4 14.4
ancestral crRNA - VOC D80A 89.5 0.8 15.5 22.8
VOC crRNA - VOC D80A 100.0 3.6 113.0 79.7
ancestral crRNA - ancestral L452R 100.0 13.9 95.5 98.9
VOC crRNA - ancestral L452R 99.4 8.6 79.0 76.5
ancestral crRNA - VOC L452R 101.1 0.7 4.0 12.0
VOC crRNA - VOC L452R 100.0 4.2 88.7 80.6
ancestral crRNA - ancestral S477N + T478K 100.0 8.5 108.2 101.9
VOC crRNA - ancestral S477N + T478K 66.3 0.7 8.9 14.2
ancestral crRNA - VOC S477N + T478K 105.6 69.4 107.1 120.6
VOC crRNA - VOC S477N + T478K 100.0 81.4 94.7 94.3
WT R377A N378A R973A
100
50
% cleaved product vs. WT PM

**Figure S13 | Related to Figure 5. A-C.** Heatmaps of percentage of cleaved product for each LbuCas13a variant when activated with a SARS-CoV-2 short RNA fragment, compared to wild-type (WT) LbuCas13a with a perfect match RNA (PM), and using a truncated crRNA. Values assessed after 1 hour incubation using 10 nM of target RNA. crRNA and target were either designed for the ancestral/Wuhan strain or to a given variant-of-concern (VOC) strain for the S gene as follows: beta (D80A), delta (L452R), omicron (S477N+T478K region). The different designs are **A.** SNP occurs at position 7 relative to crRNA; **B.** enzyme is primed with a mismatch at position 19 in all cases and SNP occurs at position 7 relative to crRNA; **C.** enzyme is primed with a mismatch at position 6 in all cases and SNP occurs at position 7 relative to crRNA.

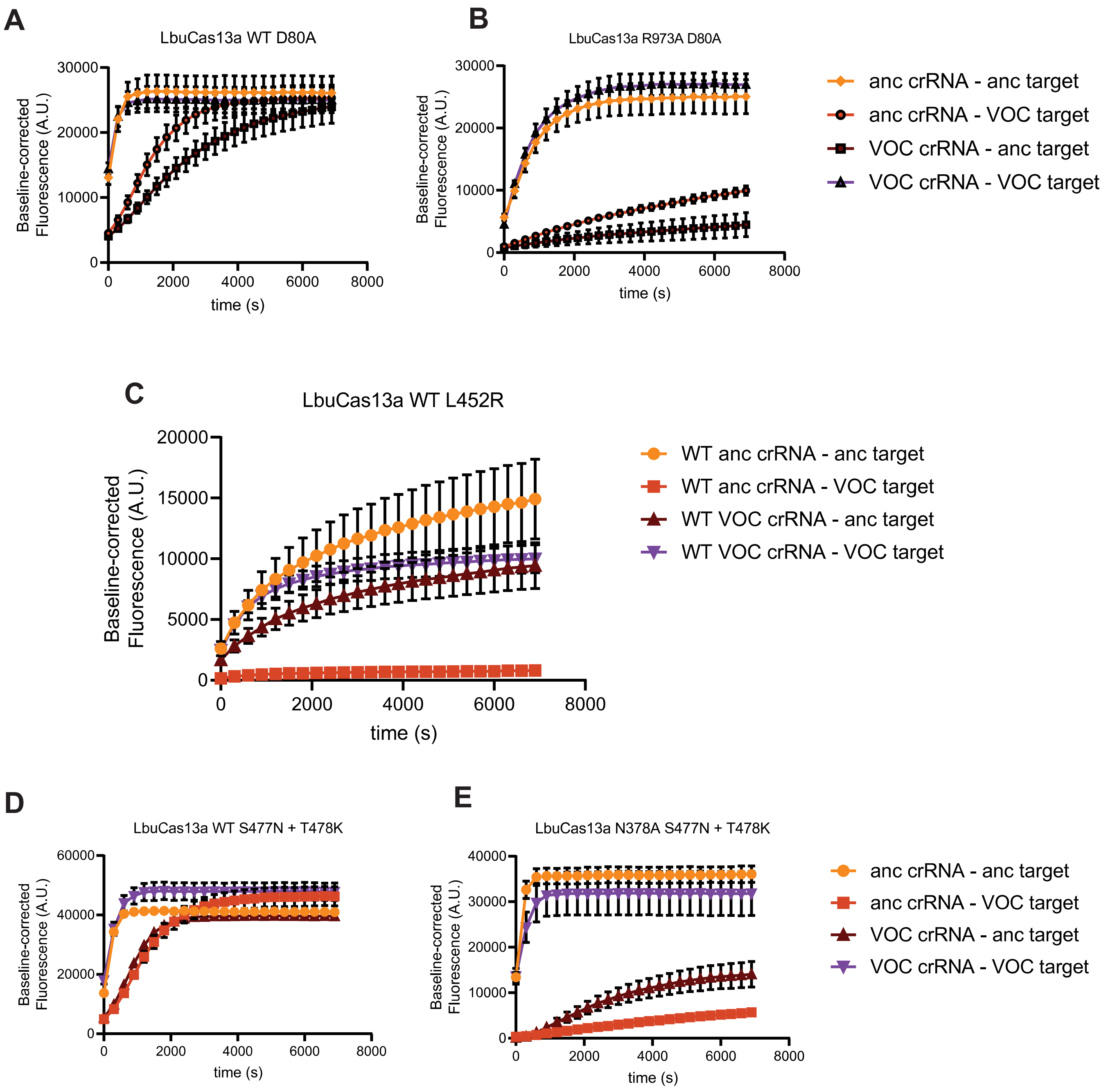


**Figure S14 | Related to Figure 5.** LbuCas13a fluorescent reporter cleavage time-course for the detection of SARS-CoV-2 variants by pre-amplification of viral RNA extracts and T7-transcription. **A.** Time-course fluorescent signal from WT when detecting a SARS-CoV-2

spike:D80S strain SNP or ancestral strain with a crRNA specific for the viral variant or the ancestral virus. **B.** Time-course fluorescent signal from LbuCas13a$^{R973A}$ when detecting a SARS-CoV-2 spike:D80S strain SNP or ancestral strain with a crRNA specific for the viral variant or the ancestral virus. **C.** Time-course fluorescent signal from WT when detecting a SARS-CoV-2 spike:L452R strain SNP or ancestral strain with a crRNA specific for the viral variant or the ancestral virus. **D.** Time-course fluorescent signal from WT when detecting a SARS-CoV-2 spike: S477N+T478K strain SNPs or ancestral strain with a crRNA specific for the viral variant or the ancestral virus. **E.** Time-course fluorescent signal from LbuCas13a$^{N378A}$ when detecting a SARS-CoV-2 spike:S477N+T478K strain SNPs or ancestral strain with a crRNA specific for the viral variant or the ancestral virus.

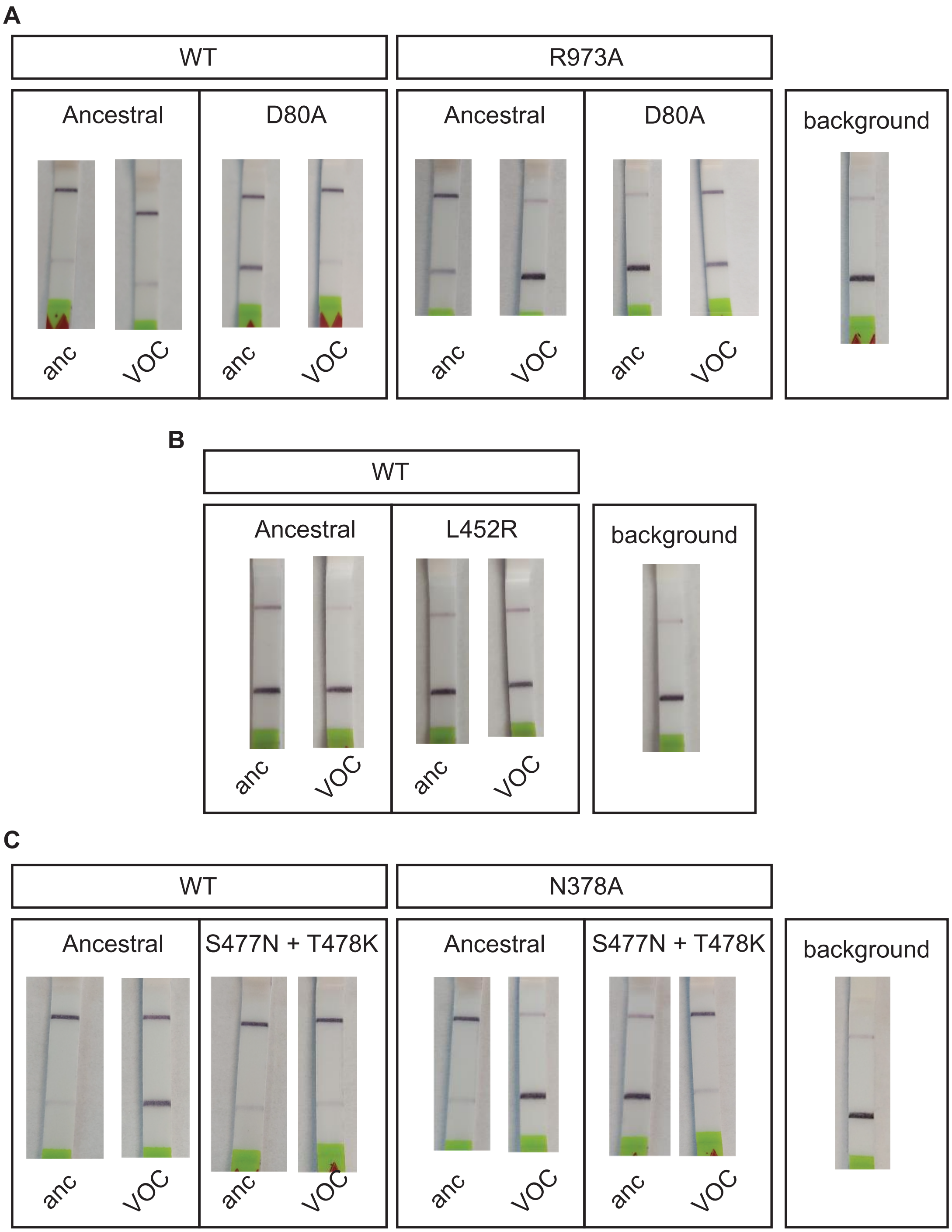


**Figure S15 | Related to Figure 5.** Lateral flow readout after 1 hour incubation of LbuCas13a reporter cleavage for the detection of SARS-CoV-2 variants by pre-amplification of viral RNA extracts and T7-transcription. **A.** Lateral flow detection after one-hour reaction from WT and

LbuCas13a$^{R973A}$ when detecting a SARS-CoV-2 spike:D80S strain SNP (VOC target) or ancestral strain (anc. target) with a crRNA specific for the viral variant (VOC crRNA) or the ancestral virus (anc. crRNA). **B.** Lateral flow detection after one-hour reaction from WT when detecting a SARS-CoV-2 spike:L452R strain SNP (VOC target) or ancestral strain (anc. target) with a crRNA specific for the viral variant (VOC crRNA) or the ancestral virus (anc. crRNA). **C.** Lateral flow detection after one-hour reaction from WT and LbuCas13a$^{N378A}$ when detecting a SARS-CoV-2 spike:S477N+T478K strain SNP (VOC target) or ancestral strain (anc. target) with a crRNA specific for the viral variant (VOC crRNA) or the ancestral virus (anc. crRNA).

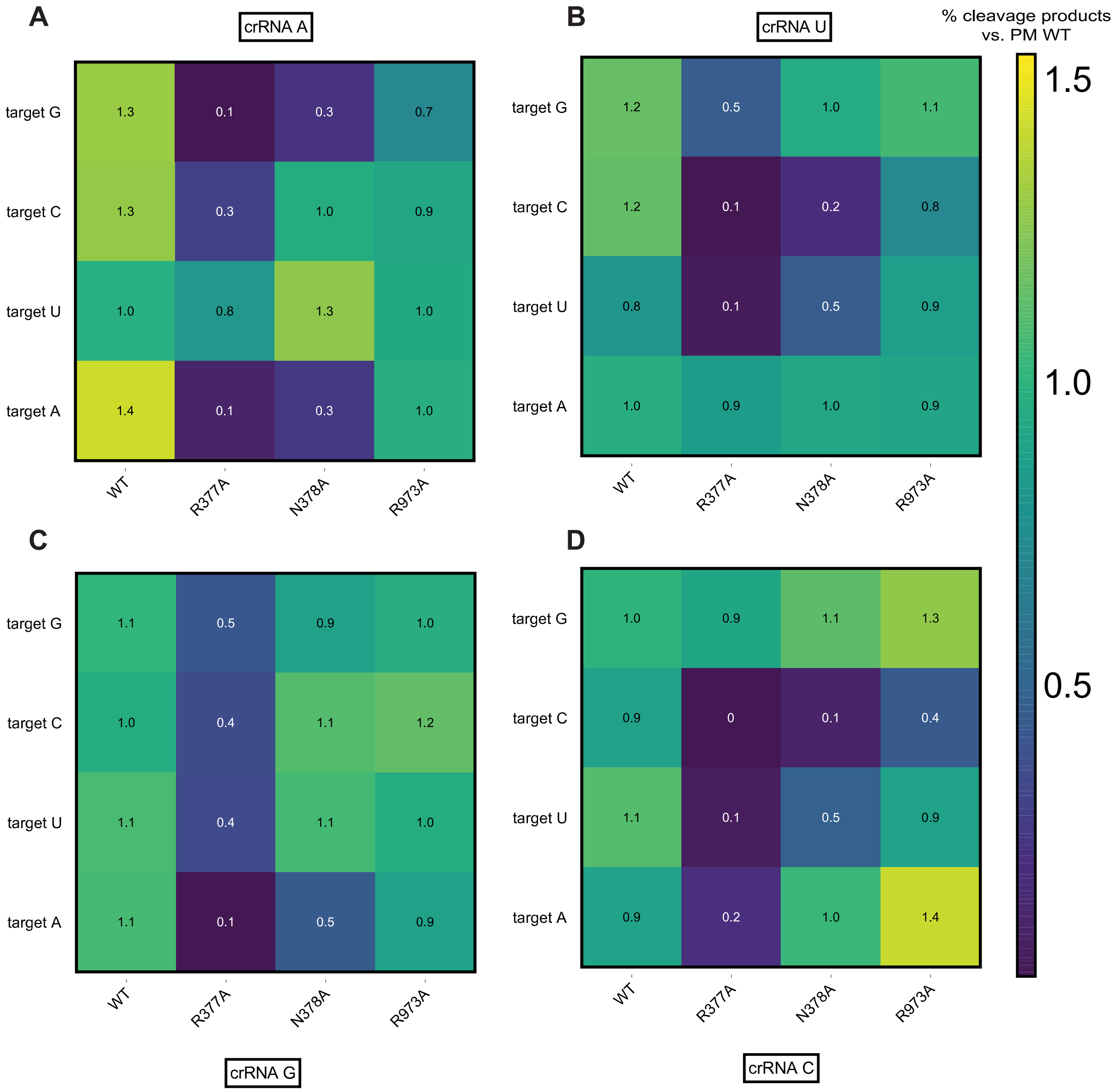


**Figure S16 | Related to Figure 6. The type of mismatch and local sequence context modulates Cas13-mismatch tolerance.** Heatmaps of percentage of cleaved product for each LbuCas13a variant and target RNA, compared to wild-type (WT) LbuCas13a with a perfect match RNA (PM), and using a truncated crRNA. Each heatmap indicates the nucleotide composition of the target RNA at position 7 of the crRNA:target duplex, with different nucleotide base pairing combinations depending on the crRNA as follows: **A.** crRNA has an adenine at this position; **B.**

crRNA has a uridine at this position; **C.** crRNA has a guanidine at this position; **D.** crRNA has a cytidine at this position.

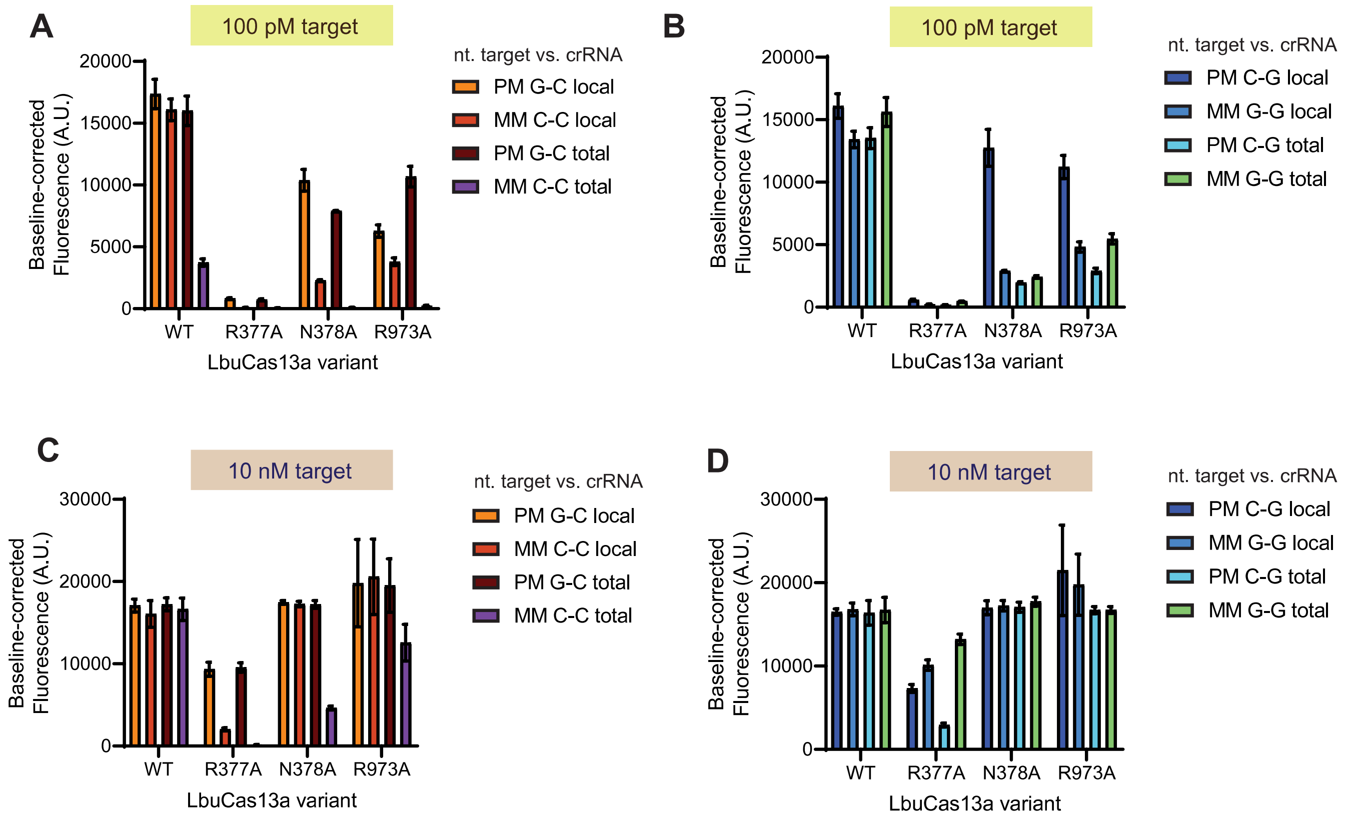


**Figure S17 | Related to Figure 6.** Comparison of one hour end-point fluorescence signal from LbuCas13a and variants with different mismatched base pairs and target RNA concentrations, for the derived RNA sequences with different GC content. **A.** Fluorescent signal for LbuCas13a variants loaded with a crRNA with C at position 7 of the crRNA:target and with 100 pM of different target RNAs that contain a mismatch (MM) or not (PM). **B.** Fluorescent signal for LbuCas13a variants loaded with a crRNA with G at position 7 of the crRNA:target and with 100 pM of target RNA. **C.** Fluorescent signal for LbuCas13a variants loaded with a crRNA with C at position 7 of the crRNA:target and with 10 nM of different target RNAs that contain a mismatch (MM) or not (PM). **D.** Fluorescent signal for LbuCas13a variants loaded with a crRNA with G at position 7 of the crRNA:target and with 10 nM of target RNA. Local refers to the derived target RNAs with high GC content around the mismatched position, keeping the original 25% GC content and total refers to derived RNAs with raised GC content to 50% but keeping the original GC content around the mismatched position, as shown in **Figure 6B-C**.
